# Electrophysiological Markers of Fairness and Selfishness Revealed by a Combination of Dictator and Ultimatum Games

**DOI:** 10.1101/2021.08.23.457310

**Authors:** Ali M. Miraghaie, Alessandro E. P. Villa, Reza Khosrowabadi, Hamidreza Pouretemad, Mohammad A. Mazaheri, Alessandra Lintas

**Affiliations:** Neuroheuristic Research Group, Faculty of Business and Economics, University of Lausanne, CH 1015 Lausanne, Switzerland & Faculty of Psychology, Shahid Beheshti University, Deneshjou Bouleverd, Evin Tehran, Iran; Neuroheuristic Research Group, Faculty of Business and Economics, University of Lausanne, CH 1015 Lausanne, Switzerland; Institute for Cognitive and Brain Science, Shahid Beheshti University, Tehran, Iran; Dept. of Clinical & Health Psychology, Shahid Beheshti University, Tehran, Iran; Neuroheuristic Research Group & LABEX, Faculty of Business and Economics, University of Lausanne, CH 1015 Lausanne, Switzerland

**Keywords:** Cooperation, EEG, N1, P2, MFN, Late positive potential, Spiteful punishment, Costly punishment

## Abstract

Event Related Potentials (ERPs) were recorded from 39 participants who played the role of Allocators in a Dictator Game (DG) and Responders in an Ultimatum Game (UG). Most participants expressed very low levels of altruistic decision making, and two homogeneous groups could be identified, one formed by fair (*N* = 10) individuals and another by selfish (*N* = 8) individuals. At fronto-central cortical sites, the ERP early negativity (N1) was reduced in selfish participants with a latency about 10 ms earlier than in fair participants. In fair DG players, the features of the subsequent positive wave P2 suggested that more cognitive resources were required when they allocated the least gains to the other party. P2 latency and amplitude in the selfish group supported the hypothesis that these participants tended to maximize their profit, as expected by a rational *Homo economicus*. During UG, we observed that a medial frontal negativity (MFN) occurred earlier and with greater amplitude when selfish participants rejected less favorable endowment shares. In this case, all players received zero payoffs, which showed that MFN in selfish participants was associated with a spiteful punishment. At posterior-parietal sites we found that the greater the selfishness, the greater the amplitude of the late positive component (LPC). Our results bring new evidence to the existence of specific somatic markers associated with the activation of distinct cerebral circuits by the evaluation of fair and unfair proposals in participants characterized by different expressions of perceived fairness, thus suggesting that particular brain dynamics could be associated with moral decisions.

## 1 Introduction

The maximization of individual’s own gain is considered to represent the main behavioral drive during an economic transaction. However, the balance of individuals’ self-interest and acceptance of others’ benefit is a common observation based on culturally shaped moral phenomena such as fairness and altruism (Rachlin, 2002; Fehr and Rockenbach, 2004; Altman, 2005; Henrich et al., 2010). Unrelated individuals try to ensure fairness to others in daily life, and hope to get fair treatment and cooperation in return (Rand and Nowak, 2013; Kocher et al., 2012). Strong reciprocity model, as one of the strategies for maintaining and urging people to commit to the norms, has many practical and effective use in the light of cooperation (Gintis et al., 2003; Stallen and Sanfey, 2013) and people act differently in diverse societies to apply it (Herrmann et al., 2008). Cooperation induced by the sacrifice of one’s own self-interest in punishing violations of the social norm defines the so-called altruistic punishment (Fehr and Gächter, 2002; Fowler, 2005; Du and Chang, 2015; Balafoutas et al., 2016). A different kind of punishment, so-called spiteful punishment, appears when an individual spends personal resources to punish one or several other individuals who behave against his presumed interest within the context of collective norms and rules (Jensen, 2010; Marlowe et al., 2011; Brañas-Garza et al., 2014; Yamagishi et al., 2017).

The Ultimatum Game (UG) has been one of the most prolific experimental tasks aimed at investigating the nature of human fairness over the last decades (Güth et al., 1982). A typical UG involves two players, the Proposer, who offers how to share an endowment in two parts, and the Responder, who can either accept the offer to share it accordingly or to reject it with both players receiving a zero payoff. Empirical studies have demonstrated that Proposers typically offered approximately 40% (fair offers) of the total amount at stake, with Responders being more likely to reject 20% or less of positive, albeit low (unfair) offers (Camerer, 2003; Sanfey et al., 2003). Rejection of an unfair offer at UG might be triggered by cooperative and popular motivations or it can be considered to be an expression of a costly punishment aimed at enforcing the social norm of fairness (Fehr and Schmidt, 1999; Henrich et al., 2006; Fehr and Fischbacher, 2004; Hewig et al., 2011). This argument has been used to support the strong reciprocity model of the evolution of cooperation (Bear and Rand, 2016; Righi and Takács, 2018; Chen et al., 2019), but competitive and spiteful motivation associated with less prosocial attitude have also been suggested (Kirchsteiger, 1994; Brañas-Garza et al., 2014). The Dictator Game (DG) task (Kahneman et al., 1986; Artinger et al., 2010) is basically a single player game because the Proposer (called the Allocator in DG) determines at his own will whether to send a fraction (ranging from nothing to all) of the initial endowment to the Responder (called the Recipient in DG). The Recipient plays a passive role and has no influence over the outcome of the game. It has been observed that the vast majority of Allocators give something to the Recipients, usually about 20% of the amount at stake, despite the fact that Allocators have no reason to give up some of the money (Artinger et al., 2010; Camerer, 2003).

Contactless monetary transactions are rapidly increasing in all socio-cultural environments and in the absence of effective face-to-face negotiations it seems crucial to gain a better understanding of decision-making processes. The observation that individuals characterized by well-defined impairments in decision making, as they tend to take decisions not advantageous for their personal lives and for their social status, may also be characterized by biophysical measurements associated with brain activity led to define the Somatic Markers Hypothesis (Bechara et al., 1994; Bechara and Damasio, 2005; Rilling and Sanfey, 2011; Villa et al., 2012). The observation of such somatic markers is important to investigate the neural basis of decision making processes. In UG and DG, the key question is searching of where and how a “decision” is taken in a recursive way. Functional magnetic resonance imaging (fMRI) has been mainly used to investigate the localization of brain activity associated with the violations of social norms and expectations in decision-making tasks (Sanfey et al., 2003; Spicer et al., 2007). The search of ‘how’ requires a fine grain temporal resolution (in the order of 1 ms time scale), which is a feature of electroencephalography (EEG) by means of recording brain activity in a non invasive way with external electrodes placed over many standard locations of the scalp (Boksem and De Cremer, 2010; Wu et al., 2011a; Luo et al., 2014; Yu et al., 2015; Mesrobian et al., 2015; Peterburs et al., 2017; Cui et al., 2019).

Transient electric potentials associated in time with a sensory or mental occurrence are termed event-related potentials (ERPs) (Picton et al., 2000). A fundamental prediction of this model is that multiple occurrences of the same event evoke a similar pattern of activity in the same cerebral circuits. Hence, the usual analysis of ERPs requires to average the activity triggered by multiple repetitions of the same event. Among the ERP wave components observed in cognitive studies, we introduce those that have been associated to experimental protocols of interest with respect to our study. The most common triggering events are the stimulus onset corresponding, in UG, to the time of the decision (acceptance/rejection) taken by the Responder following the presentation of the offer by the Proposer and, in DG, to the time of the decision (which endowment share offered to the Recipient) taken by the Allocator. In the chronological order, we introduce N1 (peaking near 130 ms after stimulus onset), P2 (or P200, peaking at 220 ms), medial-frontal negativity (MFN, peaking at 310 ms), and a late positive component (LPC, extending between 400 and 650 ms). In several experimental paradigms, ERPs can be stimulus-locked at the time of presentation of the feedback information on the decision made. This is not the case in the current study and ERPs triggered by the onset of the feedback phase are not further discussed.

In a UG protocol, the amplitude of frontal N1 was modulated by the facial attractiveness of the player (Ma and Hu, 2015; Weiss et al., 2020) and following observed nonverbal social interactions (friendly vs. nonfriendly) with Proposers (Moore et al., 2021). Parietal N1 amplitude was reported in association with the process of emotional stimuli regardless of trustworthiness of the player (Mei et al., 2020). In UG, it was observed that an increase in P2 is larger in response to high-value over low-value offers (Weiss et al., 2020), in participants informed by the opponent’s low social status (Hu et al., 2014). A similar observation was reported in participants who feel gratification in response to rewarding results but also by seeing the opponent player’s bad luck (Falco et al., 2019). The MFN amplitude was larger following unfair as compared to fair offers (Polezzi et al., 2008; Boksem and De Cremer, 2010; Wu et al., 2011a; Alexopoulos et al., 2012; Mesrobian et al., 2015). In DG, MFN is observed in Recipient’s ERP after receiving disadvantageous inequitable allocations (Li et al., 2020), but it is also dependent on the recipients’ social status and altruistic tendency (Sun et al., 2015; Cui et al., 2019; Mayer et al., 2019). A late positive component (LPC) observed at latencies of 450-650 ms over central and parietal areas was sometimes termed late positive potential and often regarded as a sustained P300 response to be associated with emotion-driven decision making (Ito et al., 1998; Wu et al., 2011b; Cui et al., 2019). In UG, the P300/LPC amplitude was larger in conditions with higher risk-tendency, such as following responses to a fair offer made by an unknown Proposer in comparison with the same payoff offered by a socially known and close individual (Polezzi et al., 2010; San Martín, 2012; Yu et al., 2015; Lintas et al., 2017; Harris et al., 2020). This amplitude also tended to be larger and localized more frontally after rejection of unfair offers (Mesrobian et al., 2015) and after receiving fair over unfair offers (Xu et al., 2020).

An unanswered question from previous studies, which usually pooled participants in decision-making tasks in one group, is whether somatic markers are different in individuals with different profiles of fairness towards accepting/rejecting wretched offers in UG. This question has been investigated in this study with an original design combining DG and UG. The cooperation is tested with the same participant playing the roles of Allocator in DG and Responder in UG, thus allowing to separate individuals’ profiles following an index of *altruism* determined during UG and an index of *selfishness* determined during DG. The experimental prediction is that the combination of these games will reveal the existence of different groups of individuals characterized by differences in selfishness and altruism. To our knowledge, this is the first study to carry out a systematic comparison of ERPs following the behavior of the same participants performing both DG and UG neuroeconomic games. The fronto-central N1 wave component is expected to be associated with top-down prefrontal control of decision making, meaning that N1 should be different in groups, if any, clearly separated by selfish and altruistic behaviors. Rewarding choices are expected to evoke larger P2 in participants who feel gratification, meaning that P2 is expected to be larger in altruistic and fair participants rather than in selfish and conceit participants. Being associated with disadvantageous inequitable offers and with altruistic tendency, MFN is expected to be larger in altruistic participants. The amplitude of posterior-parietal LPC is associated with motivation and subsequent allocation of attention, meaning that it is expected to be large in selfish Allocators playing DG and in altruistic Responders playing UG. An additional careful analysis of N1, P2, MFN, and LPC latencies for distinct behavioral groups is expected to point out interacting factors, if any, associated with the outcome of decision making.

## 2 Method

### 2.1 Participants

An *a priori* power analysis was conducted to compute the required sample size on the basis of repeated measures and within-between interaction (Faul et al., 2007). We planned 6 measurements per value (taken by three experienced users at two adjacent electrode sites) and two factors, each one with two levels (i.e. the accept/reject response and the UG/DG game), using a desired effect size of 0.33 (to represent a medium effect) with a power of 0.95 (1-*β* error probability). The power analysis indicated we need a sample size of 44 to achieve this result. Forty-four volunteer healthy young male adults were recruited via posted announcement at the university campus of the Shahid Beheshti University of Tehran (Iran), but only 39/44 completed the protocol. Then, our participant count is restricted to *N* = 39 volunteers aged between 18 and 32 years old (M=22.5, SEM=0.5). This sample size corresponds to a power of 0.92. All participants followed an academic education (8 undergraduate, 22 postgraduate and 9 PhD students) and all were right-handed with normal or corrected-to-normal vision. The research protocol was approved by the mandatory Ethics Committees requested by Shahid Beheshti University and with written informed consent from all participants, in accordance with the latest version of the Declaration of Helsinki (World Medical Association., 2000). Prior to the experimental session, all participants were interviewed to asses that they were not reporting any neurological or neuropsychiatric diseases and not taking any psychoactive medication. The whole experiment lasted between 2 and 2 and half hours. The participants were compensated for their time with a fixed amount of cash (200,000 IRR, Iranian Rials, approximately corresponding to 4 USD). In addition, the participants received performance-based cash rewards (comprised between 200,000 and 800,000 IRR) corresponding to the cumulative outcome of the UG and DG earned at the end of the experimental session.

### 2.2 Experimental procedure

The participants were seated comfortably in a sound-attenuated, electrically shielded, and dimly lit cabin with the instruction to maintain their gaze on a white fixation cross at the center of a 32-inch computer screen at a viewing distance of 100 cm. At the begin of the recording session, the EEG was recorded during 2 min while the participants kept the eyes closed and during 2 min while they fixated a cross on the center of the computer screen. The participants were randomly assigned in two groups, those who played UG at first and those who played DG at first. Each game started with 10 trials of practice. After completing the first game, a short break (5-10 min) was set to allow the participant to get acquainted with the rules of the second game.

Each game was composed of 5 blocks with 48 trials each (240 trials for each game) and the participants took a break after each block. Each trial started with a fixation cross appearing for 800-1200 ms. Then, a circular pie chart representing the endowment at stake divided in two colored pieces, was presented with the diameter of 8.0 visual degree centered on a 32-inch computer screen at a 100 cm viewing distance. The pie chart appeared on the screen for 1200 ms. The red piece of the pie corresponded always to the participant’s payoff of the endowment share. The participant had up to 3000 ms to make the decision either to accept the endowment subdivision represented by the pie chart —by pressing the ‘K’ letter of the computer keyboard with the medium finger of the right hand– or to reject it —by pressing the ‘H’ letter with the index finger of the same hand. During this interval, a schematic drawing appeared on the computer screen with a ‘thumb up’ emoji above a letter ‘K’ on the right and a ‘thumb down’ emoji above a letter ‘H’ on the left. After making the decision, the players fixated the cross for 800-1200 ms.

At the end of each trial, the outcome of the game showing the payoff earned by the participant and his opponent, was presented on the computer screen for 2000 ms. The stimulus presentation and data acquisition were performed using PsychLab and EEG8 amplifiers (Precision Instruments, Glastonbury, Somerset, BA6 8AQ, UK). The statistical analyses were performed using the R language and environment for statistical computing (R Core Team, 2021; Venables and Ripley, 2002).

### 2.3 Dictator Game

In the Dictator Game of this study, the Allocator/Proposer could not impose *a priori* the amount to be divided with the other player (the Recipient/Responder), but he had to refuse or agree with a randomly selected allocation appearing on the computer screen. As in a usual Dictator Game, the Recipient/Responder was completely passive and had to accept whatever amount was imposed by the Allocator/Proposer. Both players received zero payoff if the Allocator/Proposer refused the randomly selected allocation. All participants played the role of Allocator/Proposer and they were informed that the Recipient/Responder was an anonymous player located elsewhere. The participants played 240 trials subdivided into five blocks of 48 trials. The allocations were randomly selected from among five ([Allocator/Proposer]: [Recipient/Responder]) possibilities {(50:50), (60:40), (70:30), (80:20), (90:10)} —a fraction (80:20) meaning 80% of the amount allocated to the participant and 20% allocated to the other player. A single block of 48 trials was not necessarily balanced for the five possible fractions, but the complete sequence of 240 (5 × 48) trials was uniformly distributed, such that the sequence contained the same number of trials for each pre-determined allocation. In this version of DG, the performance-based cash reward of the Participant (always playing the role of Allocator/Proposer) was determined by the average payoff of all accepted proposals. Figure 1 presents a schematic illustration of the time course of two DG trials.

**Figure 1:**
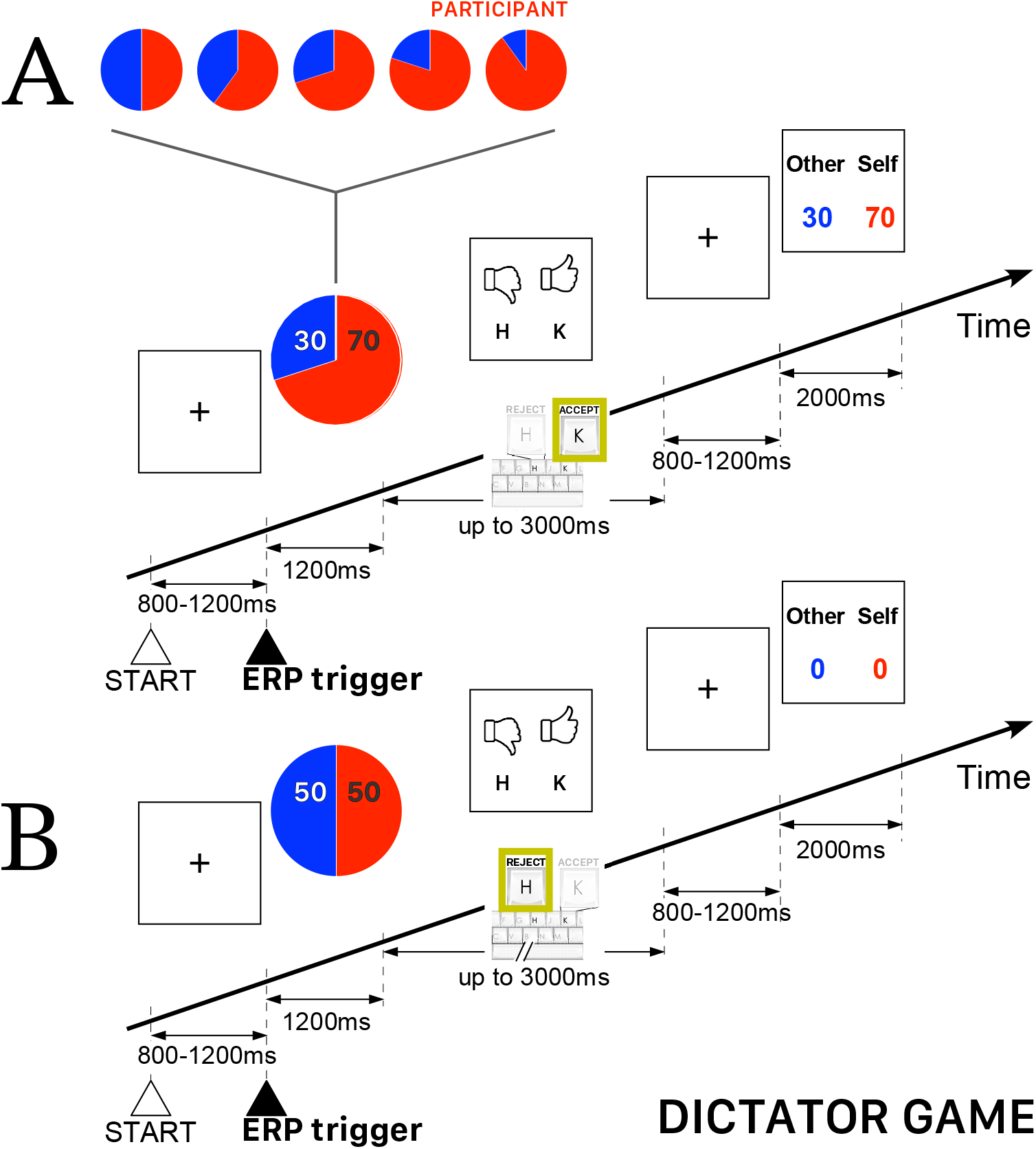
Schematic illustration of the time course of the Dictator Game task. The participant always played the role of Allocator/Proposer with the possibility to accept or refuse an allocation randomly presented on the computer screen. The red piece of the pie corresponds to participant’s payoff. **A.** Example of a trial when the Allocator/Proposer agreed with the randomly proposed allocation —in this example, 70% for the participant and 30% for the other player. **B.** Example of a trial when both players ended with a zero payoff because the Allocator/Proposer refused the randomly proposed allocation (50:50).

### 2.4 Ultimatum Game

All participants played the role of Responder, who could either accept or reject the offers appearing on the computer screen. The participants were informed that the Proposer was an anonymous player located elsewhere. In case of acceptance, each player received a payoff according to the proposed split, otherwise both players did not receive anything. The overall sequence of 240 trials was subdivided into five blocks of 48 trials with five possibilities of (Proposer:Responder) endowment shares {(50:50), (60:40), (70:30), (80:20), (90:10)} —a fraction (80:20) meaning 80% of the endowment paid to the Proposer (i.e., the opponent player) and 20% paid to the Participant, who always played the role of Responder. The trials were uniformly randomly distributed such that the complete sequence contained the same number of trials for each pre-determined endowment share. The performance-based cash reward of the participant was determined, as usually in UG, by the total payoff of all accepted endowment shares. Figure 2 presents a schematic illustration of the time course of two UG trials.

**Figure 2:**
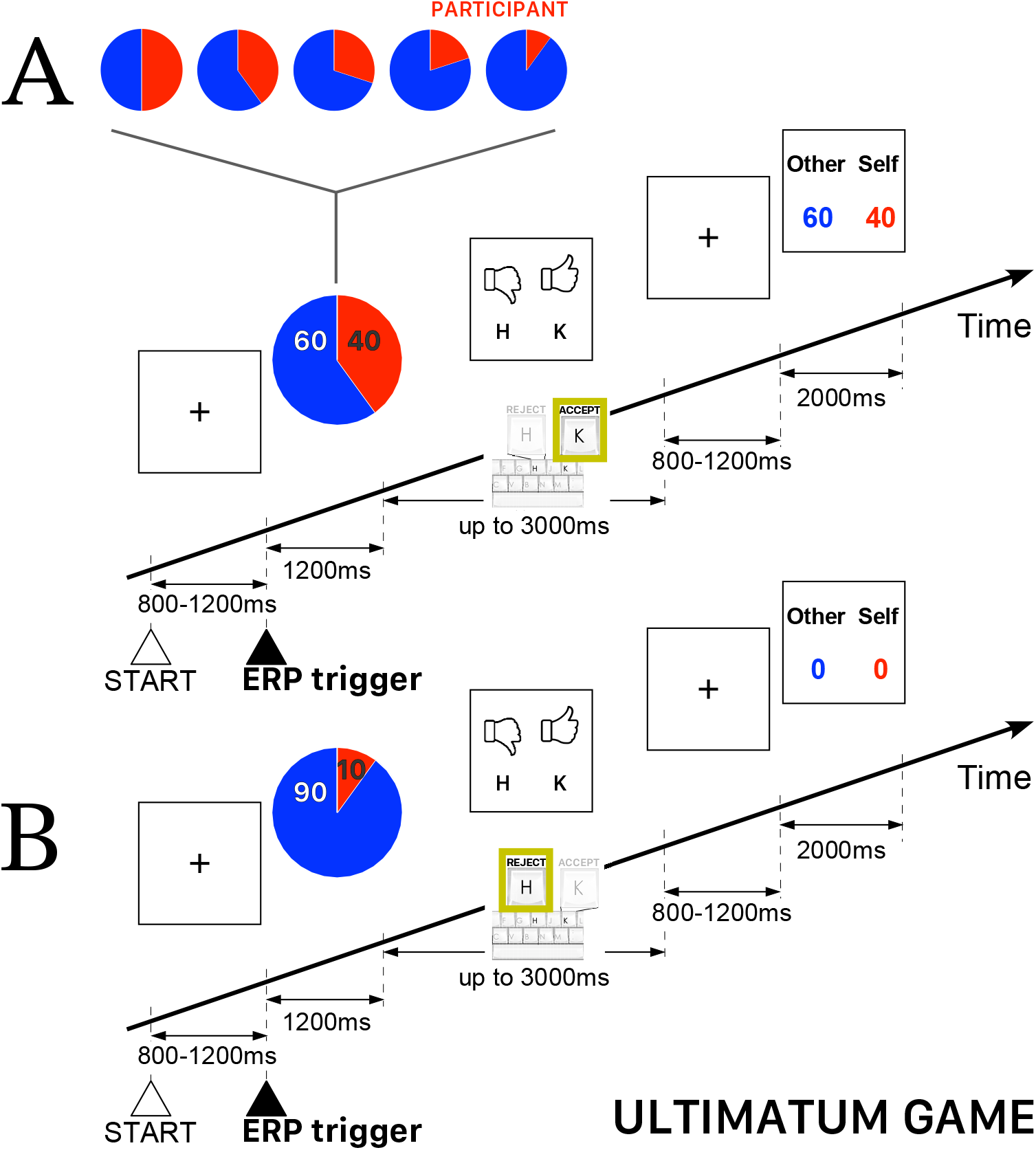
Schematic illustration of the time course of the Ultimatum Game task. The Participant always played the role of Responder and could either accept or reject the endowment share randomly presented on the computer screen. The red piece of the pie corresponds to participant’s payoff. **A.** Example of a trial when a Participant accepted the randomly proposed share—in this example, 60% for the other player and 40% for the Participant playing the role of Responder. **B.** Example of a trial when a Participant rejected the randomly proposed share—in this example, 90% for the the other player and 10% for the Participant—thus ending with a zero payoff for both players.

### 2.5 EEG recording and ERP processing

The electroencephalogram (EEG) of each Participant was recorded using 32 scalp dry titanium/titanium nitride (Ti/TiN) electrodes, mounted on a headcap (international 10/20 layout) referenced to the left mastoid bone. Electrophysiological signals were sampled at 1024 *Hz* and filtered with a band-pass of 0.1 – 200 Hz, with electrode impedances kept near 5 *k*Ω for all recordings. The analysis was performed using the EEGLAB_v12 (Delorme and Makeig, 2004)) and ERPLAB_v7 toolboxes (Lopez-Calderon and Luck, 2014)) of MATLAB 2017b (The Mathworks, Inc., Natick, MA, USA). Raw data were preprocessed with a band-pass IIR Butterworth filter from 0.1 to 32 Hz (–36*dB*/octave roll off). Blink, saccade and eyelid artifact components were corrected or set to zero, based on their respective shape and topography after using Independent Component Analysis (ICA) (Jung et al., 2000; Plöchl et al., 2012). Epochs time-locked to markers were obtained after off-line segmentation of the continuous EEG. The epochs were defined for the interval between −200 through 800 ms around the trigger and corrected to baseline 200 ms prior to marker. Noisy data were rejected after further visual inspection of the epochs for contamination by muscular or electrode artifacts. We analyzed four ERP components defined by the early negative peak near 110–160 ms after the trigger (N1), followed by a positive wave peaking in the 180–240 ms interval post trigger (P2), a negative component observed in the 260-400 ms interval (MFN), and the most positive peak observed in the 430-630 ms interval (LPC). For each participant and at each electrode site, measurements of each peak latency and amplitude, if peaks of wave components were clearly identified, were taken by three independent trained experimenters. In this way, we could perform repeated measurements (three at maximum) for each wave component at each electrode site.

### 2.6 Statistical analyses

The statistical software R v4.0.5 (R Core Team, 2021). was used for all the analyses with packages effectsize (Makowski et al., 2019), rstatix (Kassambara, 2020) and sjstats (Lüdecke, 2021). In general, the grouped values were reported as (median, Mean ±SEM). The null hypothesis of homoscedasticity (i.e., equal variances) in the data samples was tested with the Levene’s test. Unpaired comparisons between groups were performed by Student *t*-test if variances were equal and by Welch *t*-test, otherwise. For multiple comparisons, the *p*-value of *t*-test was adjusted following Bonferroni correction. The factorial analysis was performed using linear mixed effects models (with package lmerTest; Kuznetsova et al., 2017) and goodness of fit was assessed by F-statistics based on Satterthwaite’s method for denominator degrees-of-freedom and *F*-statistic). The generalized eta squared 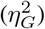 was used to estimate the effect size of main and interaction factors (*negligible* if 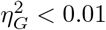; *small* if 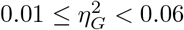; *medium* if 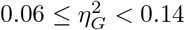; *large* if 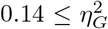). For independent two-samples comparisons, the effect size of *t*-test was assessed by Cohen’s *d (negligible* if *d* < 0.2; *small* if 0.2 ≤ *d* < 0.5; *medium* if 0.5 ≤ *d* < 0.8; *large* if 0.8 ≤ d).

## 3 Results

### 3.1 Behavioral analysis

On average (M±SEM), the participants to the Dictator Game completed 237.3±0.7 out of 240 trials. This difference is due to the fact that, in a few trials, some participants did not take any decision (to agree or refuse the proposed allocation) within the maximum interval of 3000 ms. The valid DG trials were subdivided in three sets: (*i*) *DG selfish behavior* including the trials when the participant playing the role of Allocator/Proposer agreed with the most favorable proposals of allocations for himself {(90:10), (80:20)} (i.e. 90% or 80% of the amount in favor of the participant) and refused the least favorable amounts {(60:40), (50:50)}; (*ii*) *DG neutral behavior* including all trials (both agreed and refused) with an allocation (70:30); (*iii*) *DG fair behavior* including the trials when the participant playing the role of Allocator/Proposer agreed with the least favorable allocations for himself {(60:40), (50:50)} (i.e. 60% or 50% of the amount in favor of the participant) and refused the most favorable allocations {(90:10), (80:20)}. Hence, on the basis of the number (Nb.) of trials belonging to these three sets, we computed the index *DG_selfishness_* (in the range [−1,+ 1]), such that the higher the index the more selfish the behavior, defined as follows:

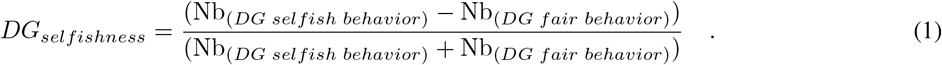

In the Ultimatum Game, all participants played the role of Responder and completed 238.1±0.4 trials. The behavioral analysis during UG was based on the subdivision of valid trials in the next three sets: (*i*) *UG altruistic behavior* including the trials when the Responder accepted the least favorable offers for himself {(90:10), (80:20)} (i.e. 10% or 20% of the share for himself) and rejected the equitable splits {(60:40), (50:50)}; (*ii*) *UG neutral behavior* including all trials (both accepted and rejected) offering an endowment share (70:30); (*iii*) *UG conceit behavior* including the trials when the Responder accepted the equitable offers {(60:40), (50:50)} and rejected the least favorable offers {(90:10), (80:20)}. Notice that in this version of UG, the equitable splits {(60:40), (50:50)} corresponded also to the most favorable payoffs that the participant could expect to receive. In a way similar to DG, we computed an index termed *UG_altruism_* (in the range [−1,+1]) based on the the number of trials in each set, such that the higher the index the more altruistic (i.e., the less conceit) the behavior, defined as follows:

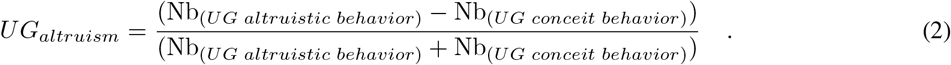

On the basis of both indices *DG_selfishness_* and U*G_altruism_*, we performed an agglomerative hierarchical clustering using the function `hclust` set with a metric Euclidean distance and the complete-linkage option of the standard `stats` package of R (R Core Team, 2021). At the begin, the procedure considered the possibility of 39 clusters, i.e. one cluster for each participant. Then, the algorithm analyzed iteratively the outcome of the clustering such that the number of clusters decreased at each step and and the procedure finally stabilized and stopped with four clusters, labeled ‘GrpS’, ‘GrpB’, ‘GrpC’, ‘GrpF’. The procedure was repeated with different random seeds at the initialization. The outcome of repeating the procedure was that the attribution of three individuals to groups ‘GrpS’ or ‘GrpB’ depended on the initial random seed. Then, we defined an additional cluster labeled ‘GrpA’ formed by those three individuals.

Table 1 reports the behavioral features of the clusters of participants and cluster ‘GrpA’ was composed by three participants whose behavior was intermediate between ‘GrpS’ and ‘GrpB’. Participants who expressed a conceit behavior during UG but a low level of selfishness during DG formed the stable cluster of fair participants (GrpF). Participants with a very high value of *DG_selfishness_* and the lowest level of *UG_altruism_* formed the stable cluster of selfish participants (GrpS). Cluster GrpB corresponded to participants with a medium–low *UG_altruism_* index and medium–high *DG_selfishness_* index. GrpB participants could be considered a group of more altruistic/less conceit participants within the context of this study. The most heterogeneous group of participants (GrpC) was characterized by variable *UG_altruism_* and medium–low *DG_selfishness_*. Figure 3 shows the scatterplot of the 39 participants distributed on a 2D feature space defined by their corresponding values of *UG_altruism_* and *DG_selfishness_*.

**Figure 3:**
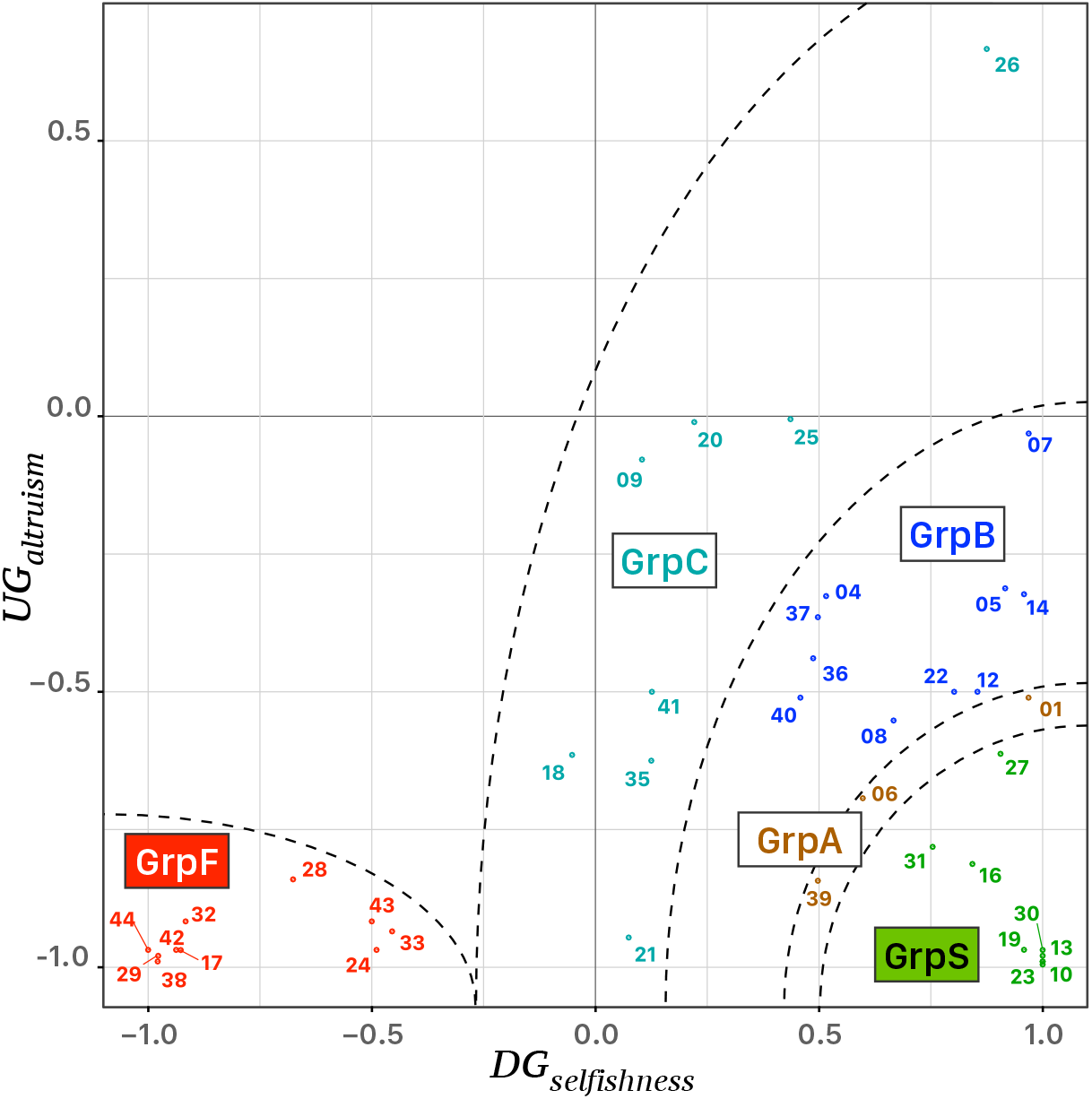
Scattergram of the behavioral analysis of 39 participants, identified by their identification tag, distributed in a 2D feature space defined by the corresponding values of *UG_attruism_* and *DG_selfishness_*. Five clusters of colored points were identified on the bases of an agglomerative hierarchical clustering procedure, i.e. ‘GrpS’ (green), ‘GrpA’ (brown), ‘GrpB’ (blue), ‘GrpC’ (turquoise), and ‘GrpF’ (red). The dashed lines correspond to ideal separatrix lines in the feature space.

**Table 1:** Relative frequencies (median, Mean±SEM) of the behavioral responses to the Dictator and Ultimatum Game and corresponding behavioral indices *DG_selfishness_* and *UG_altruism_* of the five groups of participants sorted after an agglomerative hierarchical clustering procedure described in the text.

| | $N = 39$ | 'GrpS'<br>($n = 8$) | 'GrpA'<br>($n = 3$) | 'GrpB'<br>($n = 10$) | 'GrpC'<br>($n = 8$) | 'GrpF'<br>($n = 10$) |
| --- | --- | --- | --- | --- | --- | --- |
| *Dictator Game* |  |  |  |  |  |  |
| Selfish behavior (%) |  | 79.2 | 65.0 | 69.4 | 45.0 | 3.1 |
| | | 77.4±1.3 | 67.9±5.7 | 68.5±2.6 | 49.5±4.1 | 8.6±2.9 |
| Fair behavior (%) |  | 0.8 | 16.4 | 10.6 | 34.9 | 77.0 |
| | | 2.7±1.3 | 12.6±5.8 | 11.5±2.7 | 30.4±4.1 | 71.5±2.9 |
| $DG_{selfishness}$ | | 0.98 | 0.60 | 0.73 | 0.13 | -0.92 |
| | | 0.93±0.03 | 0.69±0.14 | 0.71±0.07 | 0.24±0.10 | -0.79±0.07 |
| *Ultimatum Game* |  |  |  |  |  |  |
| Altruistic behavior (%) |  | 1.3 | 12.2 | 23.9 | 28.4 | 1.3 |
| | | 4.5±2.0 | 12.7±3.8 | 24.5±1.9 | 29.4±7.2 | 2.2±0.6 |
| Conceit behavior (%) |  | 78.7 | 67.5 | 56.1 | 51.5 | 78.6 |
| | | 75.4±2.0 | 67.1±3.9 | 55.4±1.9 | 50.5±7.2 | 77.8±0.51 |
| $UG_{altruism}$ | | -0.97 | -0.69 | -0.40 | -0.29 | -0.97 |
| | | -0.89±0.05 | -0.68±0.10 | -0.39±0.05 | -0.26±0.18 | -0.95±0.01 |

### 3.2 Reaction Times

Reaction times were measured between the time of the presentation of the allocation proposal in DG (presentation of the endowment share in UG) and the time of pressing the letter key on the computer keyboard. In either game, separately, the RTs for all 39 participants were faster when the participants agreed *vs*. refused the allocation during DG (median RT 381.0, 492.0 ± 4.9 ms *vs*. 408.0, 514.2 ± 5.1 ms; Student *t*(9093) = −3.146, *p* = .002, *d* = −0.07) or accepted *vs*. rejected the proposed endowment share during UG (median RT 403.0, 500.3 ± 4.3 ms vs. 426.5, 534.1 ± 6.4 ms; Student *t*(8961) = −4.516, *p* < .001, *d* = −0.10). Despite these differences, RTs were distributed as usually, i.e. a positively skewed distribution with a long tail which reflected occasional slow responses. These long-tailed distributions of RTs introduced a bias that was amplified by the differences between single participants. For this reason, to account within and between participants variance of raw RTs, a mixed model ANOVA with repeated measurements did not reveal the main effect of response choice (DG: *F*(1, 76) = 2.519, *p* = .117, 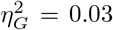; UG: *F*(1, 74) = 1.694, *p* = .197, 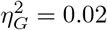). Instead of raw RTs, *z*-scores of RTs computed for each participant and each game separately were considered further for factorial analysis using mixed models with repeated (i.e., single trials) measurements. With *z*-scores of RTs the effect of factor *response* was significant in each game (DG: *F*(1, 76) = 14.47, *p* < .001, 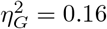; UG: *F*(1, 74) = 24.62, *p* < .001, 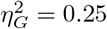).

We focused on less conceit/more altruistic (GrpB), fair (GrpF), and selfish (GrpS) participants, which corresponded to the three most homogenous groups revealed by the behavioral analysis. For each game separately, we run an analysis with factors *response* (2 levels: Agreed/Accepted and Refused/Rejected in DG/UG) and *group* (3 levels: GrpB, GrpF and GrpS) with repeated measurements. A linear mixed model fit by maximum likelihood was performed due to the presence of unbalanced samples (Table 2). A strong main effect of Response was observed in both games, with reaction times faster when the participants agreed with the allocation or accepted the proposed endowment share irrespective of the participants’ group.

**Table 2:** Reaction times. The table reports measurements of the reaction times (RTs in ms) (median, mean ±SEM values and number of trials *n*) when participants agree (or disagree) with the proposed allocation during the Dictator Game, and when they accept (or reject) the proposal of endowment share during the Ultimatum Game. Statistics of Levene’s test for heteroscedascity (homogeneity of variances test) and ANOVA-like tables for a linear mixed model fit by maximum likelihood with corresponding main factors and interaction are reported for *z*-scores of RTs computes for each participant separately.

| Factor |  | Dictator Game: Allocator |  | Ultimatum Game: Responder |  |
| --- | --- | --- | --- | --- | --- |
|  |  | Agreed | Refused | Accepted | Rejected |
| Response choice |  |  |  |  |  |
| Participants' Group |  |  |  |  |  |
| GrpB (N = 10) |  | n = 855 | n = 1032 | n = 1481 | n = 395 |
|  |  | 355.0 | 373.5 | 390.0 | 409.0 |
|  |  | 423.5±9.1 | 464.5±9.5 | 502.1±8.7 | 556.2±20.3 |
| (z-score) |  | (-0.031±0.029) | (0.024±0.029) | (-0.027±0.022) | (0.110±0.052) |
| GrpF (N = 10) |  | n = 763 | n = 1077 | n = 890 | n = 922 |
|  |  | 398.0 | 432.0 | 400.0 | 430.0 |
|  |  | 515.1±12.5 | 543.6±11.1 | 470.9±9.5 | 520.9±11.4 |
| (z-score) |  | (-0.066±0.033) | (0.047±0.027) | (-0.064±0.028) | (0.065±0.031) |
| GrpS (N = 8) |  | n = 769 | n = 738 | n = 724 | n = 692 |
|  |  | 391.0 | 415.0 | 440.0 | 426.5 |
|  |  | 488.3±11.0 | 521.8±12.1 | 521.7±11.8 | 512.6±13.0 |
| (z-score) |  | (-0.064±0.030) | (0.066±0.035) | (-0.032±0.031) | (0.036±0.036) |
| Homogeneity of variances |  |  |  |  |  |
| Levene's test: |  | F(5, 50) = 0.921, p = .475 |  | F(5, 50) = 9.008, p < .001 |  |
| Anova-like table |  |  |  |  |  |
| Effect: | Response | F(1, 50) = 12.41, p < .001 |  | F(1, 50) = 23.76, p < .001 |  |
|  | Group | F(1, 50) = 0.225, p = .800 |  | F(2, 50) = 2.688, p = .078 |  |
|  | Response × Group | F(1, 50) = 0.537, p = .588 |  | F(2, 50) = 3.108, p = .053 |  |
| Participants' Behavior |  |  |  |  |  |
|  |  | Fair trials |  | Altruistic trials |  |
|  |  | n = 862 | n = 1099 | n = 622 | n = 77 |
|  |  | 395.5 | 429.0 | 463.0 | 298.0 |
|  |  | 518.6±12.3 | 533.2±10.2 | 597.8±16.5 | 625.3±78.5 |
| (z-score) |  | (-0.047±0.035) | (0.020±0.029) | (0.242±0.048) | (0.573±0.217) |
|  |  | Selfish trials |  | Conceit trials |  |
|  |  | n = 1525 | n = 1748 | n = 2473 | n = 1932 |
|  |  | 364.0 | 390.0 | 393.0 | 428.5 |
|  |  | 448.3±6.9 | 494.2±8.0 | 472.6±5.7 | 521.0±7.7 |
| (z-score) |  | (-0.070±0.023) | (0.063±0.026) | (-0.145±0.016) | (0.047±0.023) |
| Homogeneity of variances |  |  |  |  |  |
| Levene's test: |  | F(3, 88) = 1.150, p = .334 |  | F(3, 89) = 18.36, p < .001 |  |
| Anova-like table |  |  |  |  |  |
| Effect: | Response | F(1, 88) = 0.196, p = .659 |  | F(1, 89) = 4.111, p = .045 |  |
|  | Behavior | F(1, 88) = 1.537, p = .218 |  | F(1, 89) = 23.53, p < .001 |  |
|  | Response × Behavior | F(1, 88) = 0.312, p = .578 |  | F(1, 89) = 1.018, p = .316 |  |

During DG we categorized four sets of trials according to the behavior: (i) the *Selfish trials* with the Allocator/Proposer in ‘agreement’ with the randomly selected allocations {(90:10), (80:20)} (i.e. 90% or 80% of the share for the participant); (ii) the *Selfish trials* with the Allocator/Proposer ‘refusing’ the least favorable (in the current task design) allocations for himself, i.e. {(60:40), (50:50)}; (iii) the *Fair trials* with the Allocator/Proposer in ‘agreement’ with the allocations {(60:40), (50:50)}; (iv) the *Fair trials* with the Allocator/Proposer ‘refusing’ the most favorable allocations for himself, i.e. {(90:10), (80:20)}. Along a similar scheme, four sets of trials were categorized during UG: (i) the *Altruistic trials* when the Responder ‘accepted’ the least favorable outcomes that granted only 10% or 20% of the share to the participant (i.e., shares {(90:10), (80:20)}); (ii) the *Altruistic trials* when the Responder ‘rejected’ equitable offers (i.e., shares {(60:40), (50:50)}); (iii) the *Conceit trials* when the Responder ‘accepted’ the equitable offers (i.e., the participant received 40% or 50% of the share that are also the most advantageous payoffs in the current task design); (iv) the *Conceit trials* when the Responder ‘rejected’ the least favorable payoff. Note that during UG any trial ended without a payoff for the participant if he rejected the offer (either altruistic or conceit).

For both games, the factorial analysis (Table 2) was run with factors *response* (2 levels) and *behavior* (2 levels). During UG, we observed a very strong effect of behavior (RTs during conceit trials shorter than during altruistic trials) *F*(1, 89) = 4.111, *p* = .045, 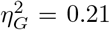) and confirmed the main effect (although weaker) of the response choice (*F*(1, 89) = 23.53, *p* < .001, 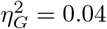). On the contrary, during DG no main effects of response and behavior were observed in groups (GrpB, GrpF, GrpS).

A multiple regression was conducted to see if the indices *DG_selfishness_* and *UG_altruism_* predicted the RTs in a specific set of trials. During DG, it was found that the level of *DG_selfishness_* alone always explained a significant amount of the variance in RTs in all sets of trials. We considered both linear and quadratic fits for raw RTs and *z*-scaled RTs as explained above, but quadratic fits always provided better results. During selfish trials (Figure 4A), irrespective of the response choice, RTs tended to be faster for participants whose *DG_selfishness_* index was high (*F*(2,53) = 17.00, *p* < .001, *R*^2^ = .391, 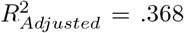). During fair trials, it is interesting to note a specular curve (Figure 4B), characterized by faster RTs for participants whose *DG_selfishness_* index was low (*F*(2,53) = 26.66, *p* < .001, *R*^2^ = .447, 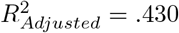). During UG, *UG_altruism_* alone explained most of the variance only in a specific set of trial. i.e. when participants rejected the proposed endowment share during conceit trials (*F*(1, 35) = 16.53, *p* < .001, *R*^2^ = .321, 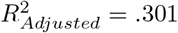).

**Figure 4:**
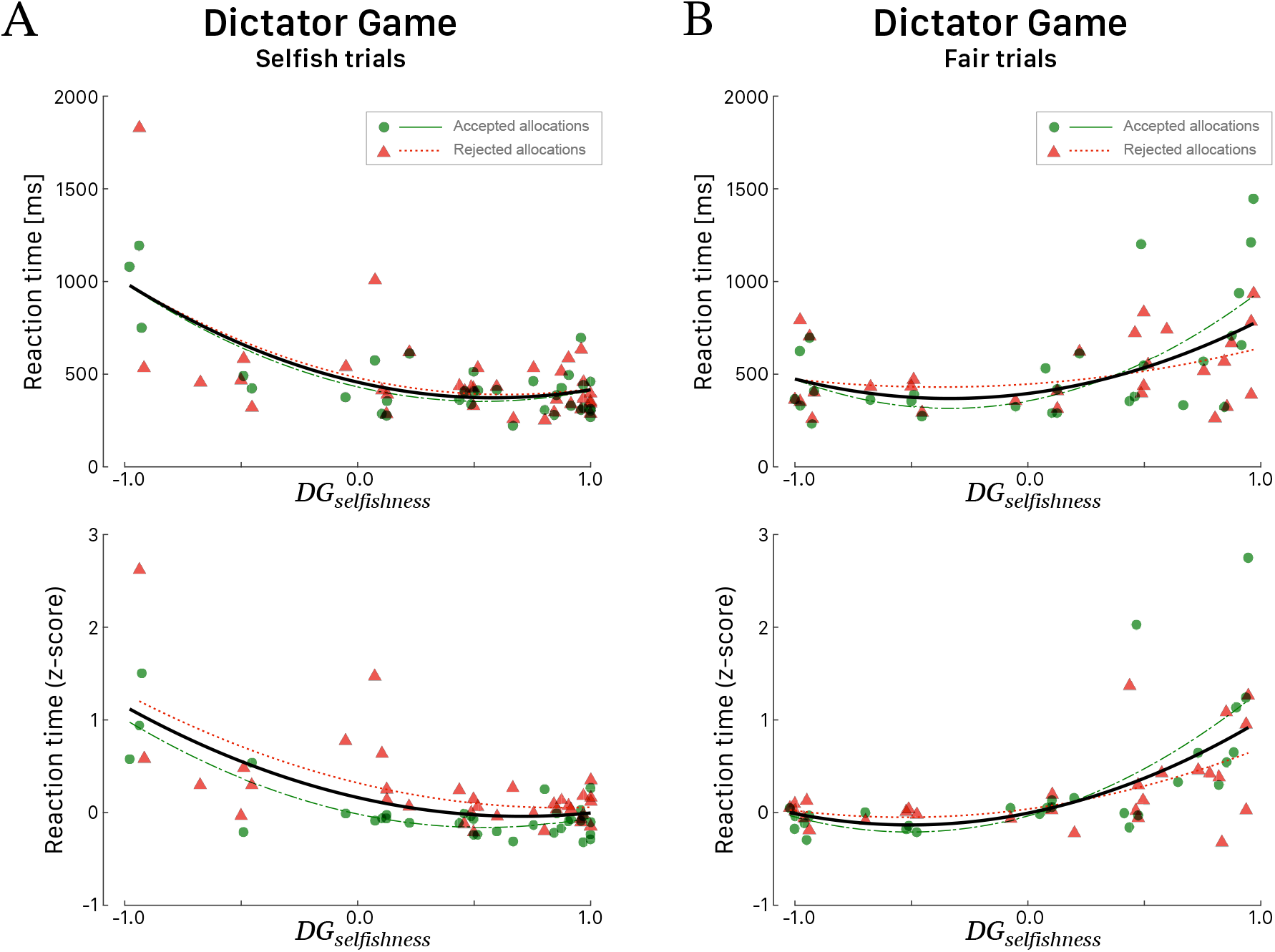
Reaction times (in ms, top panels; corresponding *z*-score in bottom panels) for all 39 participants during the Dictator Game as a function of the *DG_selfishness_* behavioral index. All participants played the role of Allocator. RTs during *Selfish trials* **(A)** and *Fair trials* **(B)**. Green dots correspond to participants’ RTs when they agreed with the proposed allocation and red triangles when they refused the allocation. Green (long dashed) and red curves correspond to quadratic fitted regressions. The black curves correspond to quadratic fits irrespective of the response choice.

### 3.3 Electrophysiological results

Figure 5 shows the grand average ERPs for all trials recorded at fronto-central (FCz and Fz) and posterior-parietal sites (Pz and CPz) triggered by the stimulus onset in both games and followed by a response (accept or reject) for all 39 participants grouped together. The visual inspection of the grand-averaged waveforms showed several components, which were generally observed in both games and at various scalp locations, but their amplitudes varied greatly as a function of the participant and of the recording site. The earliest component was a negative wave (N1) ranging in latency between 110 and 160 ms post trigger, particularly visible at fronto-central sites, and at FCz in particular. A posterior-parietal N1 was often observed, but it was tiny and noisy in the ERP of several participants, which led us to discard this wave from further quantitative analysis. This wave was followed by P2, a positive component much sharper at fronto-central sites peaking at approximately 220 ms post trigger and by a negative wave peaking within the interval 260-400 ms after trigger onset. Note that the amplitude of this negative component was deeper at fronto-central sites for both games, which led us to identify it as the medial frontal negativity (MFN). At posterior-parietal sites (Figure 5), the form and latency of this negativity were different and the component was called N2. A positive wave extending from 400 to 650 ms after stimulus onset followed MFN/N2 at all sites. This extended positivity was characterized by two successive peaks, which might be identified as the putative P3 and the late positive potential. At posterior-parietal sites, the amplitude of this positivity was larger than at other sites and the second peak was larger than the first peak. Then, the second peak at posterior-parietal sites appeared as being the most representative and was identified as the peak of the late positive component (LPC).

**Figure 5:**
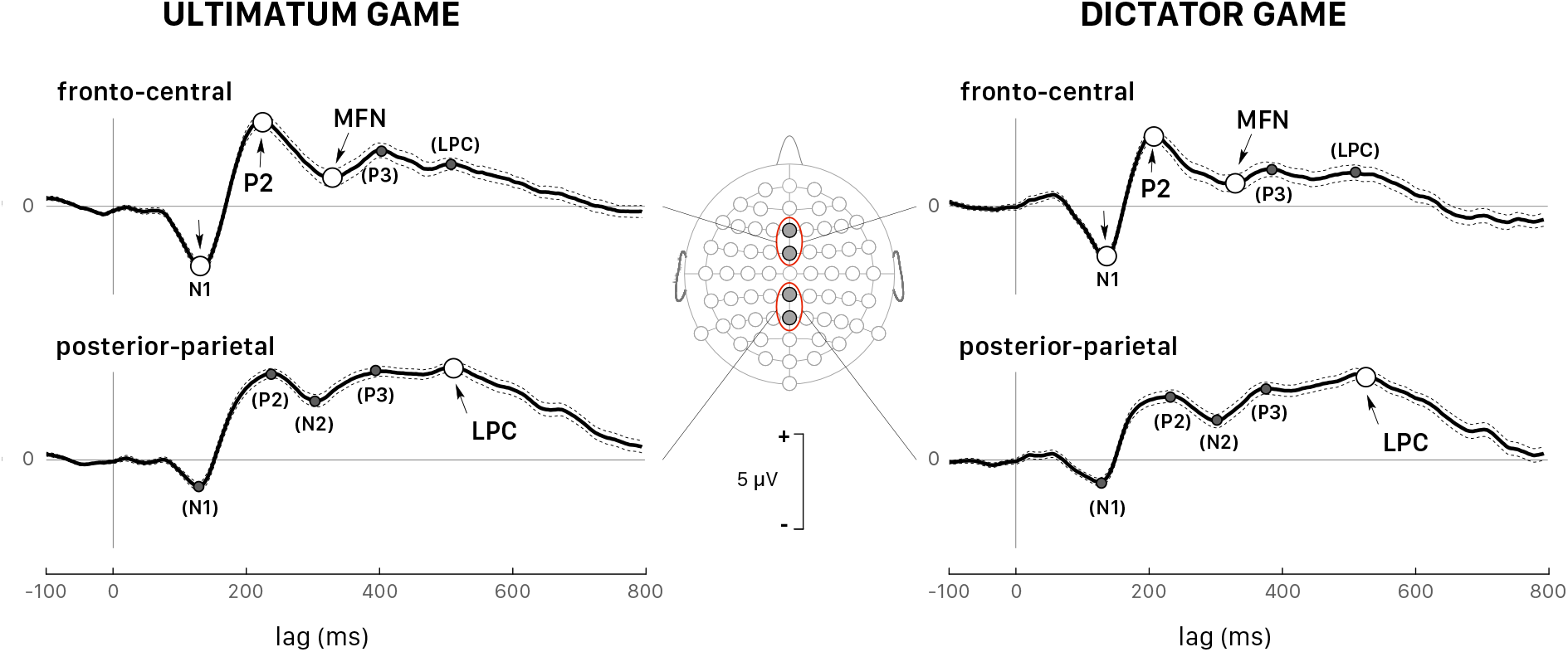
Grand average (all 39 participants and *all trials pooled together*) of the ERPs triggered by stimulus presentation during the Ultimatum Game (left panels) and the Dictator Game (right panels). The dotted lines show the confidence intervals determined by the standard error of the mean (SEM). All peaks indicated with arrows correspond to the wave components whose latency and peak were analyzed in detail: P2 and medial frontal negativity (MFN) at the fronto-central sites (Fz and FCz, upper panels) and the late positive component (LPC) at the posterior-parietal sites (Pz and CPz, lower panels). All peaks identified within parentheses indicate the wave components that were not considered in the quantitative analysis.

A comparative analysis of the grand averaged ERPs during DG and UG, with all trials pooled irrespective of payoff and participants’ decision, was carried out on peak latencies and amplitudes. The analysis was focused on less conceit/more altruistic (GrpB), fair (GrpF), and selfish (GrpS) participants, which corresponded to the three most homogenous groups revealed by the behavioral analysis. We run an analysis with factors *Game* (2 levels: UG and DG) and *Group* (3 levels: GrpB, GrpF and GrpS) with repeated measures (i.e., two electrodes for each area and three measurements at each site, as described in the Methods section). Due to the presence of unbalanced groups we performed the factor analysis with a linear mixed model fit by maximum likelihood and present the results in an Anova-like table (Table 3). Note that wave peaks were not always visible in all participants and in both games, and therefore the number of measurements (n in Table 3) for each variable was not always a regular multiple of the number of participants. For example, in group GrB with *N* =10 participants, the maximum number of measurements for a variable was *n* = 60, but it could actually be as low as *n* = 48 (e.g., N1 peak of GrpB during DG) if a peak could be uniquely identified at all electrodes.

**Table 3:** ERP wave components. The table reports measurements of the peak latency (in ms) and amplitude (in *μ*V) (median, mean ±SEM values and number of measurements *n*) of N1, P2, and MFN recorded at the fronto-central sites (Fz and FCz) and LPC recorded at the posterior-parietal sites (Pz and CPz) irrespective of the endowment share and of participants’ decision. The table reports Levene’s test for heteroscedascity (homogeneity of variances test) and type III Anova-like table for a linear mixed model fit by maximum likelihood with factors *Grp* and *Game*, and unpaired *t*test comparisons for values of only fair and selfish groups of participants.

|  |  | fronto-central N1 |  |  |  | fronto-central P2 |  |  |  |
| --- | --- | --- | --- | --- | --- | --- | --- | --- | --- |
| | | Latency [ms] | | Amplitude [$\mu V$] | | Latency [ms] | | Amplitude [$\mu V$] | |
| Factor | Game | DG: Allocator | UG: Responder | DG: Allocator | UG: Responder | DG: Allocator | UG: Responder | DG: Allocator | UG: Responder |
| Participants' Group |  |  |  |  |  |  |  |  |  |
| GrpB ($N = 10$) | | $n = 48$ | $n = 54$ | $n = 48$ | $n = 54$ | $n = 60$ | $n = 58$ | $n = 60$ | $n = 58$ |
|  |  | 139.1 | 130.8 | -4.04 | -3.94 | 222.1 | 222.7 | 5.01 | 4.72 |
| | | 139.3±1.7 | 130.0±1.2 | -4.67±0.36 | -3.92±0.20 | 217.2±2.4 | 223.4±2.0 | 4.81±0.26 | 4.77±0.14 |
| GrpF ($N = 10$) | | $n = 60$ | $n = 60$ | $n = 60$ | $n = 60$ | $n = 56$ | $n = 56$ | $n = 56$ | $n = 56$ |
|  |  | 138.4 | 140.7 | -2.83 | -3.53 | 201.3 | 226.0 | 4.40 | 4.83 |
| | | 139.2±1.2 | 141.0±2.0 | -3.14±0.28 | -3.65±0.17 | 201.8±2.5 | 227.1±3.2 | 4.40±0.26 | 4.98±0.22 |
| GrpS ($N = 8$) | | $n = 39$ | $n = 48$ | $n = 39$ | $n = 48$ | $n = 48$ | $n = 48$ | $n = 48$ | $n = 48$ |
|  |  | 128.2 | 127.8 | -1.66 | -2.33 | 205.9 | 211.3 | 3.31 | 3.74 |
| | | 128.9±2.3 | 131.9±2.0 | -1.40±0.19 | -2.66±0.30 | 207.0±3.5 | 212.5±2.4 | 4.15±0.34 | 4.12±0.32 |
| unpaired $t$-test (GrpF, GrpS) | | $t(97) = 4.357$ | $t(106) = 3.159$ | $t(97) = -4.623$ | $t(106) = -2.993$ | $t(102) = -1.239$ | $t(102) = 3.515$ | $t(102) = 0.603$ | $t(102) = 2.254$ |
| | $p$-value | $p < .001$ | $p = .012$ | $p < .001$ | $p = .018$ | $p = .218$ | $p = .004$ | $p = .548$ | $p = .156$ |
| | Cohen's $d$ | $d = 0.90$ | $d = 0.61$ | $d = -0.95$ | $d = -0.58$ | $d = -0.24$ | $d = 0.69$ | $d = 0.12$ | $d = 0.44$ |
| Homogeneity of variances |  |  |  |  |  |  |  |  |  |
| | Levene's test: | $F(5, 46) = 1.080, p = .384$ | | $F(5, 46) = 0.896, p = .492$ | | $F(5, 50) = 0.595, p = .704$ | | $F(5, 50) = 1.188, p = .329$ | |
| Anova-like table |  |  |  |  |  |  |  |  |  |
|  | Effect: |  |  |  |  |  |  |  |  |
| | Game | $F(1, 46) = 0.334, p = .566$ | | $F(1, 46) = 0.566, p = .456$ | | $F(1, 50) = 4.109, p = .048$ | | $F(1, 50) = 0.019, p = .892$ | |
| | Group | $F(2, 46) = 2.087, p = .136$ | | $F(2, 46) = 6.055, p = .005$ | | $F(2, 50) = 1.130, p = .331$ | | $F(2, 50) = 0.427, p = .655$ | |
| | Game × Group | $F(2, 46) = 1.016, p = .370$ | | $F(2, 46) = 1.334, p = .273$ | | $F(2, 50) = 1.068, p = .351$ | | $F(2, 50) = 0.083, p = .920$ | |

|  |  | fronto-central MFN |  |  |  | posterior-parietal LPC |  |  |  |
| --- | --- | --- | --- | --- | --- | --- | --- | --- | --- |
| | | Latency [ms] | | Amplitude [$\mu V$] | | Latency [ms] | | Amplitude [$\mu V$] | |
| Factor | Game | DG: Allocator | UG: Responder | DG: Allocator | UG: Responder | DG: Allocator | UG: Responder | DG: Allocator | UG: Responder |
| Participants' Group |  |  |  |  |  |  |  |  |  |
| GrpB ($N = 10$) | | $n = 60$ | $n = 60$ | $n = 60$ | $n = 60$ | $n = 60$ | $n = 60$ | $n = 60$ | $n = 60$ |
|  |  | 326.5 | 328.1 | -2.23 | 1.07 | 524.3 | 531.1 | 5.12 | 3.66 |
| | | 323.1±4.3 | 319.7±3.8 | -0.91±0.58 | 0.10±0.42 | 525.6±3.8 | 537.5±6.2 | 4.45±0.32 | 3.88±0.20 |
| GrpF ($N = 10$) | | $n = 60$ | $n = 56$ | $n = 60$ | $n = 56$ | $n = 60$ | $n = 60$ | $n = 60$ | $n = 60$ |
|  |  | 314.6 | 338.1 | 1.26 | 0.62 | 520.6 | 581.0 | 4.19 | 2.14 |
| | | 307.8±4.2 | 336.5±2.4 | 1.56±0.29 | 0.84±0.17 | 520.1±4.3 | 574.3±5.9 | 3.97±0.32 | 2.20±0.30 |
| GrpS ($N = 8$) | | $n = 48$ | $n = 42$ | $n = 48$ | $n = 42$ | $n = 48$ | $n = 48$ | $n = 48$ | $n = 48$ |
|  |  | 292.4 | 318.3 | -2.13 | 0.68 | 551.4 | 530.5 | 4.96 | 5.75 |
| | | 307.2±5.0 | 315.9±4.5 | -1.38±0.43 | -0.30±0.41 | 558.2±5.9 | 535.7±3.5 | 4.67±0.21 | 5.82±0.21 |
| unpaired $t$-test (GrpF, GrpS) | | $t(106) = 0.096$ | $t(96) = 4.309$ | $t(106) = 5.828$ | $t(96) = 2.818$ | $t(106) = -5.360$ | $t(106) = 5.293$ | $t(106) = -1.726$ | $t(106) = -9.562$ |
| | $p$-value | $p = .924$ | $p < .001$ | $p < .001$ | $p = .036$ | $p < .001$ | $p < .001$ | $p = .261$ | $p < .001$ |
| | Cohen's $d$ | $d = 0.02$ | $d = 0.88$ | $d = 1.13$ | $d = 0.58$ | $d = -1.04$ | $d = 1.02$ | $d = -0.33$ | $d = -1.85$ |
| Homogeneity of variances |  |  |  |  |  |  |  |  |  |
| | Levene's test: | $F(5, 49) = 0.861, p = .514$ | | $F(5, 49) = 2.310, p = .058$ | | $F(5, 50) = 1.503, p = .206$ | | $F(5, 50) = 0.890, p = .495$ | |
| Anova-like table |  |  |  |  |  |  |  |  |  |
|  | Effect: |  |  |  |  |  |  |  |  |
| | Game | $F(1, 49) = 1.682, p = .201$ | | $F(1, 49) = 0.416, p = .522$ | | $F(1, 50) = 1.843, p = .181$ | | $F(1, 50) = 0.510, p = .479$ | |
| | Group | $F(2, 49) = 0.561, p = .574$ | | $F(2, 49) = 2.490, p = .093$ | | $F(2, 50) = 0.969, p = .386$ | | $F(2, 50) = 4.876, p = .012$ | |
| | Game × Group | $F(2, 49) = 1.275, p = .289$ | | $F(2, 49) = 0.350, p = .706$ | | $F(2, 50) = 4.173, p = .021$ | | $F(2, 50) = 2.228, p = .118$ | |

In selfish participants (dashed green curves in Figure 6), the amplitude of N1 peak negativity was reduced in both games, which accounted for the main effect of factor *Group* (Figure 6). In fair participants (solid red curves in Figure 6), P2 peak latency was much shorter during DG (lower panel) than UG (upper panel) (*t*(110) = −6.198, *p* < .001, *d* = −1.17). During DG (lower panel), the negativity of MFN in fair participant was reduced (GrpF vs. GrpB: *t*(118) = 3.819, *p* = .001, *d* = 0.70; GrpF vs. GrpS: *t*(106) = 5.828, *p* < .001, *d* = 1.13). At posterior-parietal sites, the main effect of factor *Group* for LPC amplitude (Table 3) was determined by larger LPC wave of GrpS during UG (GrpS vs. GrpB: *t*(106) = 6.733, *p* = .001, *d* = 1.30; GrpS vs. GrpF: *t*(106) = 9.562, *p* < .001, *d* = 1.85). Note that at the fronto-central sites of selfish participants, a sustained late positive wave also appeared during UG (dashed green curves in Figure 6, upper panel). Overall, these observations highlight different brain dynamics between fair and selfish participants, but these curves were obtained with a super pool of trials, which means all trials regardless of the amount of the endowment share, and of the decision-making process with related payoff, if any. In the next two subsections, we present the most salient results on the ERP wave components for each game, separately.

**Figure 6:**
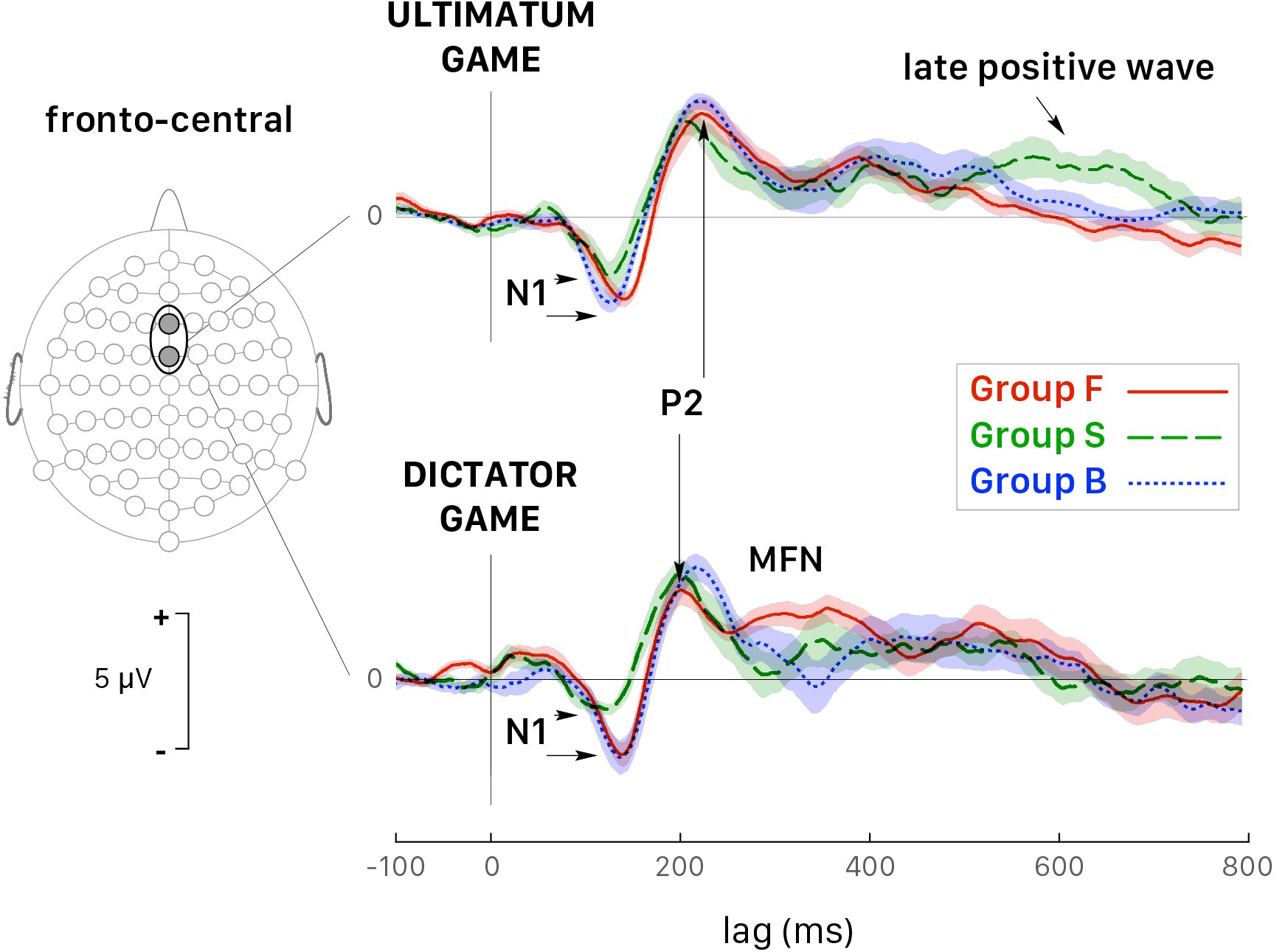
Grand average ERPs (all *trials pooled together*) recorded at fronto-central sites (Fz and FCz pooled together) during the Ultimatum Game (top panel) and the Dictator Game (bottom panel). Colored curves correspond to fair (GrpF, solid red curves), selfish (GrpS, dashed green curves) and less conceit (GrpB, dotted blue curves) participants with confidence intervals determined by the standard error of the mean (SEM). The arrows point out at several significant differences emphasized in the text. See also Table 3.

#### 3.3.1 Dictator Game

We considered separately four sets of ERP recorded trials following the same categories used for the analysis of reaction times: *Selfish trials* when the participant (i) ‘agreed’ with the highest allocations for himself and (ii) ‘disagreed’ with the least advantageous allocations; *Fair trials* when the participant (iii) ‘agreed’ with the least advantageous allocations and (iv) ‘disagreed’ with the most favorable allocations. For each participant, all single trials of one set were averaged together at each electrode site. The four sets of trials were analyzed by three observers at two electrode sites (Fz and FCz for fronto-central ERP waves and Pz and CPz for LPC). We focused our analysis on the three main groups of participants, which represented a total of 28 participants. Then, the theoretical maximum number of measurements of a kind for the whole set of four trials was *n* = 28 participants × 4 sets of trials × 2 electrodes × 3 observers = 672 for each ERP wave.

We analyzed the effect of factors *response* (2 levels: Agreed, Refused) and *behavior* (2 levels: Selfish trials, Fair trials) on latencies and amplitudes of major ERP peaks by the linear mixed model with repeated measures (Table 4). In different trials and in different participants, ERP wave peaks could not always be distinguished by all observers at all electrode sites, which ultimately represented an overall number (*n*) of measurements between 476 and 513. In order to analyze the effect of these factors independently, the table reports the values measured for the response type irrespective of the behavior and the values measured for the behavior irrespective of the response. Note that a significant main effect of participant’s *behavior* was observed on both latency and amplitude for all ERP waves (Table 4). The main effect of *response* choice was significant for MFN*_latency_* and LPC*_amplitude_*. This finding extends the differences between the groups of fair and selfish participants observed for the latency of MFN and amplitude of LPC when all trials were pooled together.

**Table 4:**
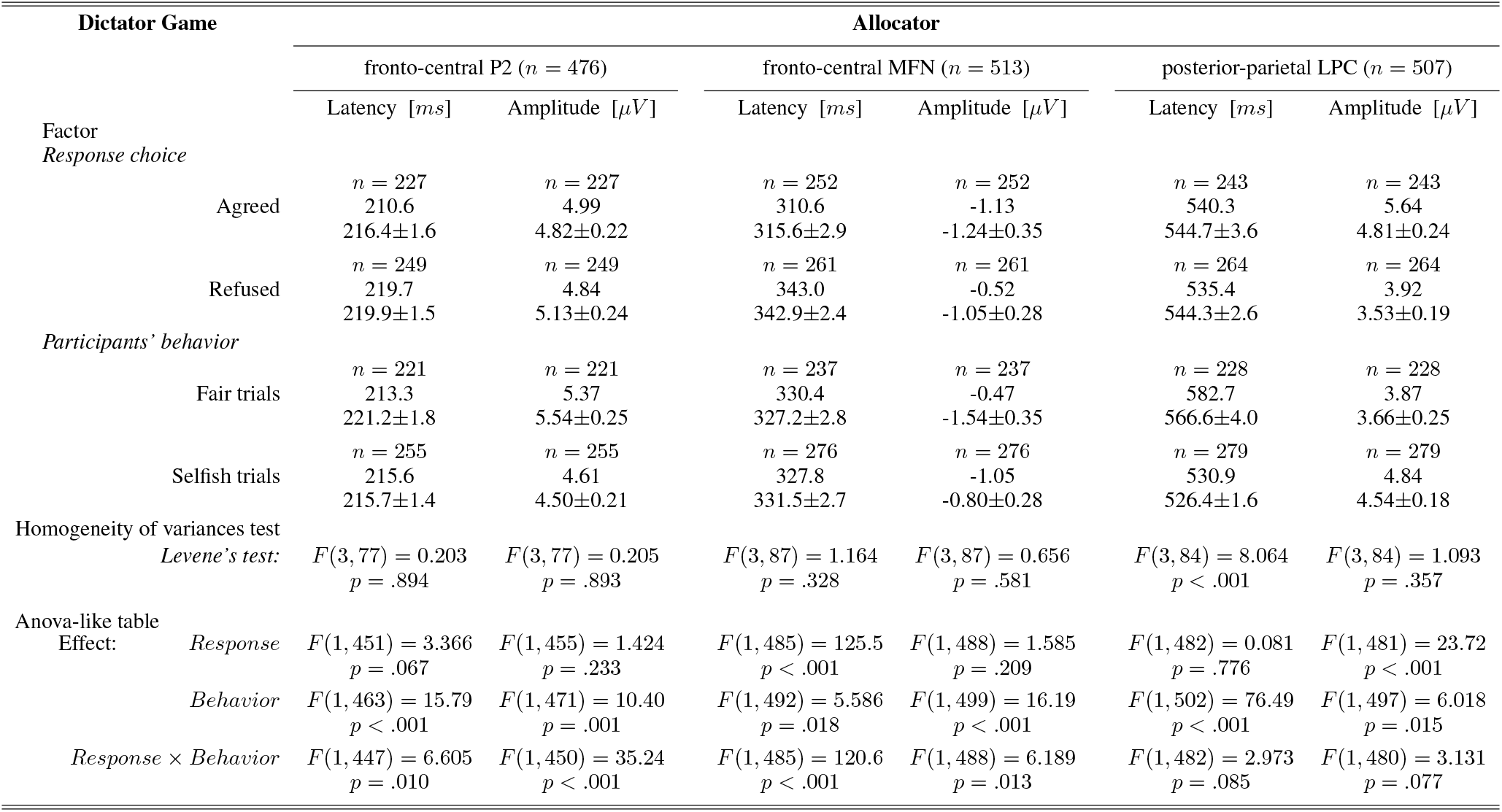
Dictator Game. All participant played the role of Allocators. The table reports measurements of each peak’s latency and amplitude (median, mean ±SEM values and number of measurements n) for trials independently sorted according to factors *response* and *behavior*. Statistics and significance are reported for Levene’s test for heteroscedascity (homogeneity of variances test) and type III Anova-like table for a linear mixed model fit by maximum likelihood.

It is interesting also to note that the interaction effects between behavior and response were statistically significant for P2 and MFN. We investigated this interaction following response choices (i.e., agreed vs. refused proposed allocations) recorded in the most representative trials for each group of participants, that were *Selfish trials* in selfish participants (GrpS) and *Fair trials* in fair participants (GrpF). Selfish participants who refused fair allocations during DG were characterized by longer P2*_latency_* (*t*(61) = 9.134, *p* < .001, *d* = 2.70) and greater P2*_amplitude_* (*t*(61) = 16.727, *p* < .001, *d* = 4.95) over refusal of selfish payoffs. This suggests that refusal of fair payoffs was a rewarding choice for selfish participants. In fair participants (GrpF), longer P2*_latency_* (*t*(88) = 3.673, *p* < .01, *d* = 0.82) was observed when they refused selfish allocations over fair ones. This suggests that refusal of selfish gains was a rewarding choice for fair participants. Accordingly, when fair participants refused selfish allocations (Figure 7A, left panel, dashed line), fronto-central P2*_amplitude_* was greater (*t*(118) = 3.43, *p* < .01, *d* = 0.63) than after acceptance of least favorable gain for themselves (Figure 7A, left panel, solid line).

**Figure 7:**
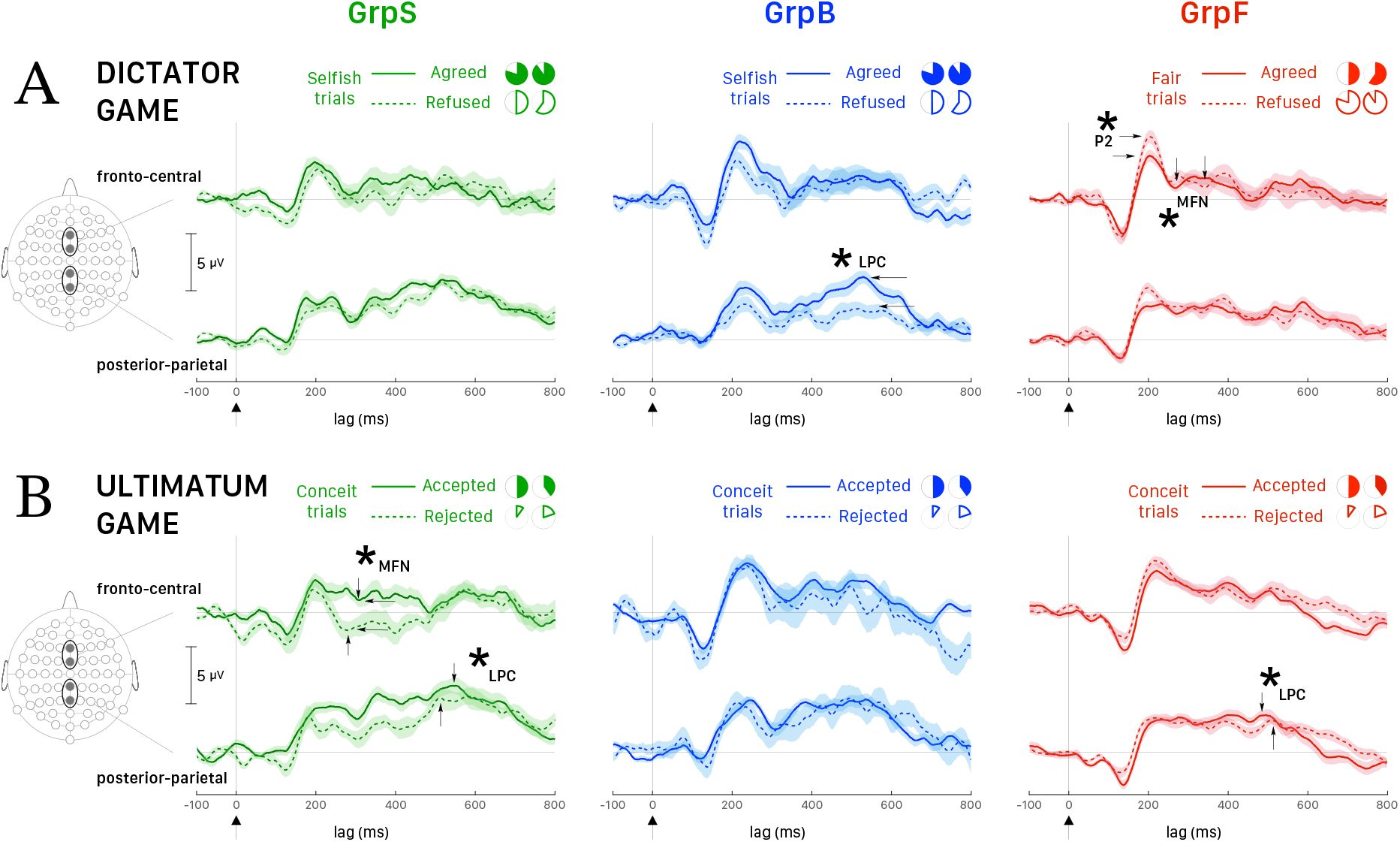
Averaged ERPs (and ±SEM bands) recorded at fronto-central sites (Fz, FCz) and posterior-parietal sites (Pz, CPz). **A.** Dictator Game: all participants played the role of Allocator/Proposer. The most representative trials for GrpS (left panel) and GrpB (middle panel) participants were *Selfish trials* and *Fair trials* for GrpF (right panel). Time zero corresponds to the presentation of the endowment share to the Allocator. Solid lines correspond to trials when the participant agreed with the proposal and dashed lines when the participant refused the proposal. **B.** Ultimatum Game: all participants played the role of Responder. The most representative trials for all groups of participants were *Conceit trials*. Time zero corresponds to the presentation of the endowment share to the Responder. Solid lines correspond to trials when the participant accepted the proposal and dashed lines when the participant rejected the proposal. Asterisks and labeled wave peaks indicate the most noticeable differences also commented in the text.

In fair participants, the depth of MFN negativity was reduced compared to the other groups (see Table 3). In GrpF, MFN*_latency_* was shorter by approximately 60 ms in fair trials (*t*(118) = −11.333, *p* < .001, *d* = −2.07) when they accepted fair (bringing the least favorable gains) over refusal of selfish (with the most advantageous payoffs) allocations (Figure 7A, left panel). In the group formed by less conceit/more altruistic participants (GrpB), MFN*_latency_* was also shorter by 60 ms (*t*(76) = −8.089, *p* < .001, *d* = −1.88) and with greater negativity (*t*(76) = 3.175, *p* < .05, *d* = 0.74) after acceptance of the least advantageous allocations during fair trials. When these participants accepted the least advantageous allocations MFN*_latency_* was also shorter by approximately 70 ms (*t*(88) = −7.567, *p* < .001, *d* = −1.69) and with greater negativity (*t*(88) = 5.029, *p* < .001, *d* = 1.12) over acceptance of selfish (with the most advantageous payoffs) allocations. Then, it appears that it is rather the proposal of the fair and least advantageous allocations that evoked larger MFN in GrpB participants.

In these participants (GrpB, the less conceit/more altruistic), the amplitude of posterior-parietal LPC (Figure 7A, middle panel) was greater in agreed (solid line) than refused (dashed line) most advantageous (selfish) trials (*t*(118) = 5.72, *p* < .001, *d* = 1.04). When these participants refused selfish allocations the LPC*_latency_* was shorter by approximately 60 ms over refusal of fair allocations (*t*(81.7) = −12.878, *p* < .001, *d* = −2.539). Among fair Allocators, the most relevant finding was a LPC*_latency_* longer by approximately 40 ms after fair proposals, irrespective of the response. Overall, these findings emphasize that different dynamics distinguished the groups of participants based on behavioral responses. Fair participants were mainly characterized by ERP markers at fronto-central sites during the early stages of decision making in our DG task, while less conceit and more altruistic participants showed most salient markers in the mid-late components of the ERPs.

#### 3.3.2 Ultimatum Game

We considered separately four sets of ERP recorded trials following the same categories used for the analysis of reaction times during UG: *Altruistic trials* when the Responder (i) ‘accepted’ the least favorable offers and (ii) ‘rejected’ equitable offers and received zero payoff; *Conceit trials* when the Responder (iii) ‘accepted’ the equitable offers (that are the most advantageous payoff in the current task design) and (iv) ‘rejected’ with the least favorable endowment shares and received zero payoff. All single trials of each set were averaged, pooled at fronto-central and posterior-parietal sites and analyzed following the same procedure used for the Dictator Game. We analyzed P2, MFN and LPC peaks, and the overall number of measurements (n) was 448, 518 and 514, respectively. Table 5 reports the latencies and amplitudes of major ERP peaks measured for the response type irrespective of the behavior and the values measured for the behavior irrespective of the response.

**Table 5:** Ultimatum Game. All participant played the role of Responders. The table reports measurements of each peak’s latency and amplitude (median, mean ±SEM values and number of measurements n) for trials independently sorted according to factors *response* and *behavior*. Same legend of Table 4.

| Ultimatum Game |  | Responder |  |  |  |  |  |
| --- | --- | --- | --- | --- | --- | --- | --- |
| | | fronto-central P2 ($n = 448$) | | fronto-central MFN ($n = 518$) | | posterior-parietal LPC ($n = 514$) | |
| Factor | | Latency [ms] | Amplitude [μV] | Latency [ms] | Amplitude [μV] | Latency [ms] | Amplitude [μV] |
| *Response choice* |  |  |  |  |  |  |  |
| Accepted | | $n = 264$ | $n = 264$ | $n = 300$ | $n = 300$ | $n = 288$ | $n = 288$ |
|  |  | 223.4 | 5.28 | 322.3 | -0.05 | 517.3 | 5.59 |
| | | 226.3±1.5 | 5.26±0.24 | 318.6±1.9 | -0.65±0.32 | 521.8±2.2 | 5.56±0.20 |
| Rejected | | $n = 184$ | $n = 184$ | $n = 218$ | $n = 218$ | $n = 226$ | $n = 226$ |
|  |  | 218.1 | 4.67 | 302.0 | -0.72 | 513.6 | 5.41 |
| | | 218.3±1.7 | 4.61±0.35 | 303.4±2.0 | -1.31±0.42 | 516.0±3.5 | 5.55±0.35 |
| *Participants' behavior* |  |  |  |  |  |  |  |
| Altruistic trials | | $n = 158$ | $n = 158$ | $n = 188$ | $n = 188$ | $n = 184$ | $n = 184$ |
|  |  | 223.0 | 5.48 | 289.8 | -3.43 | 527.9 | 6.34 |
| | | 222.9±2.4 | 5.67±0.49 | 296.7±2.3 | -2.90±0.56 | 530.0±4.2 | 6.57±0.41 |
| Conceit trials | | $n = 290$ | $n = 290$ | $n = 330$ | $n = 330$ | $n = 330$ | $n = 330$ |
|  |  | 219.9 | 4.82 | 319.8 | 0.67 | 514.4 | 4.86 |
| | | 223.1±1.2 | 4.63±0.16 | 321.0±1.6 | 0.19±0.22 | 513.3±1.9 | 4.99±0.18 |
| Homogeneity of variances test |  |  |  |  |  |  |  |
| *Levene's test:* | | $F(3, 72) = 0.409$<br>$p = .747$ | $F(3, 72) = 6.421$<br>$p < .001$ | $F(3, 83) = 1.360$<br>$p = .261$ | $F(3, 83) = 7.078$<br>$p < .001$ | $F(3, 83) = 5.632$<br>$p < .01$ | $F(3, 83) = 4.075$<br>$p < .01$ |
| Anova-like table |  |  |  |  |  |  |  |
| Effect: | *Response* | $F(1, 430) = 3.689$<br>$p = .055$ | $F(1, 441) = 1.738$<br>$p = .188$ | $F(1, 498) = 63.19$<br>$p < .001$ | $F(1, 499) = 5.624$<br>$p = .018$ | $F(1, 499) = 0.032$<br>$p = .859$ | $F(1, 499) = 5.896$<br>$p = .016$ |
| | *Behavior* | $F(1, 429) = 1.057$<br>$p = .304$ | $F(1, 439) = 5.864$<br>$p = .016$ | $F(1, 491) = 143.2$<br>$p < .001$ | $F(1, 492) = 59.10$<br>$p < .001$ | $F(1, 489) = 16.51$<br>$p < .001$ | $F(1, 489) = 36.77$<br>$p < .001$ |
| *Response × Behavior* | | $F(1, 430) = 13.80$<br>$p < .001$ | $F(1, 441) = 4.713$<br>$p = .030$ | $F(1, 497) = 2.605$<br>$p = .107$ | $F(1, 498) = 0.271$<br>$p = .603$ | $F(1, 501) = 0.282$<br>$p = .595$ | $F(1, 501) = 11.16$<br>$p = .001$ |

In the Ultimatum Game, the factors for the analysis with the linear mixed model with repeated measures were *response* (2 levels: Accepted, Rejected) and *behavior* (2 levels: Altruistic trials, Conceit trials). In the absence of main effects, the strong interaction effect observed for P2 suggested more detailed analysis. In selfish participants, it is interesting that P2*_latency_* was longer by approximately 35 ms (*t*(64) = 5.841, *p* < .001, *d* = 1.44) after acceptance of the least favorable outcomes (paying only 10% or 20% of the share to the participant) over acceptance of equitable shares (that granted 40% or 50% of the share). This comparison was weaker but held also for P2*_amplitude_*, which was greater after accepting the least favorable outcomes (*t*(53.7) = 3.204, *p* < .05, *d* = 0.80). This suggests that even acceptance of the least favorable payoffs represents a rewarding choice for selfish participants, in agreement with the hypothesis that selfish participants tend to maximize their gain at all circumstances. GrpB participants (i.e., the least conceit/the most altruistic) showed also an interesting effect on P2. In this group, the latency of P2 was longer by approximately 40 ms (*t*(70) = 4.610, *p* < .001, *d* = 1.46) and P2 amplitude was greater (*t*(12.5) = 3.422, *p* < .05, *d* = 1.25) after acceptance of least favorable over equitable shares. This supports the rationale that accepting less favorable payoffs was a rewarding choice for altruistic participants.

For MFN, significant main effects of participant’s *behavior* and *response* choice were observed on both peak latency and amplitude (Table 5). When selfish participants rejected the least favorable payoff over the acceptance of the equitable (most advantageous in this task design) offers (Figure 7B, left panel), MFN*latency* was shorter by 30 ms (*t*(94) = −5.968, *p* < .001, *d* = −1.22) and MFN negativity was greater (Welch’s *t*(93.2) = 4.003, *p* < .001, *d* = 0.82). In fair participants, after acceptance of least favorable over acceptance of equitable (most advantageous) payoffs, the latency of MFN was also shorter by approximately 30 ms (*t*(94) = −6.684, *p* < .001, *d* = −1.41) and MFN negativity was greater (Welch’s *t*(39.7) = 6.903, *p* < .001, *d* = 1.59).

The similar pattern observed for MFN at fronto-central sites between fair and selfish participants did not held any more at posterior-parietal sites. The main effect of participant’s behavior was very strong for latencies and amplitudes of LPC peak (Table 5) and the pattern for LPC was opposite between fair and selfish participants (Figure 7B). LPC*_latency_* during conceit trials, was shorter in selfish participants after rejection of least advantageous over acceptance of most advantageous (equitable) offers (Welch’s *t*(81.8) = −3.945, *p* < .001, *d* = 0.82), while it was shorter in fair participants after acceptance of most advantageous payoffs over rejection of least advantageous offers (Welch’s *t*(104.7) = 4.266, *p* < .001, *d* = 0.71).

During UG, fewer *Altruistic trials* were recorded than *Conceit trials* because participants in all groups expressed little altruism and more conceitedness. However, it is worth reporting some noticeable differences in LPC wave between the groups of participants during altruistic trials. In selfish participants, acceptance of most favorable payoffs corresponded to a shorter LPC*_latency_* by about 90 ms (Welch’s *t*(57.8) = 11.220, *p* < .001, *d* = 2.82). On the opposite, in fair participants it is the rejection of most favorable payoffs that corresponded to a shorter LPC*_latency_* by about 130 ms (Welch’s *t*(51.84) = 18.891, *p* < .001, *d* = 5.12). Participants showing less conceitedness and more altruism (GrpB) showed an intermediate pattern in comparison with selfish and fair participants. In this group, we observed shorter latency (*t*(110.7) = −5.427, *p* < .001, *d* = −1.01) and greater amplitude of LPC (*t*(108.2) = 3.584, *p*< .001, *d* = 0.67) when they accepted equitable (but also most advantageous) offers over acceptance of inequitable splits of the endowment. Overall, the analysis of ERPs during the Ultimatum Game showed selfish participants being strongly characterized at the early stages of the decision making and with a pattern of brain activity opposite to fair participants at later stages.

#### 3.3.3 Dimensional analysis of ERP peaks

The differences between the participants’ groups observed in Tables 4,5 suggest that the ERP peaks might also be correlated with the behavioral indices, which defined the participants’ groups. This analysis requires samples of data with behavioral ratings distributed throughout the values range, but the range of *UG_altruism_* was limited to values between −1 and 0 due to the bias towards conceitedness. The most significant correlations were observed with peak latencies, that are characterized by less variance than amplitudes. During fair trials of DG, fronto-central P2 peak latency correlated linearly with *DG_selfishness_* (Pearson correlation coefficient *r* = 0.63, 95% CI [0.38, 0.79], *t*(36) = 4.778, *p* < .001) when the participant agreed or refused the allocation (Figure 8A). Irrespective of the response choice, the slopes of both regression curves were similar and the smaller the selfishness, the shorter the latency of P2 peak (*F*(1, 40) = 17.45, *p* < .001, *R*^2^ = .304, 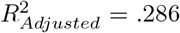). This results should be viewed taking into account the correlation between RTs and selfishness during DG (i.e., the smaller the selfishness, the shorter the RTs, Figure 4B).

**Figure 8:**
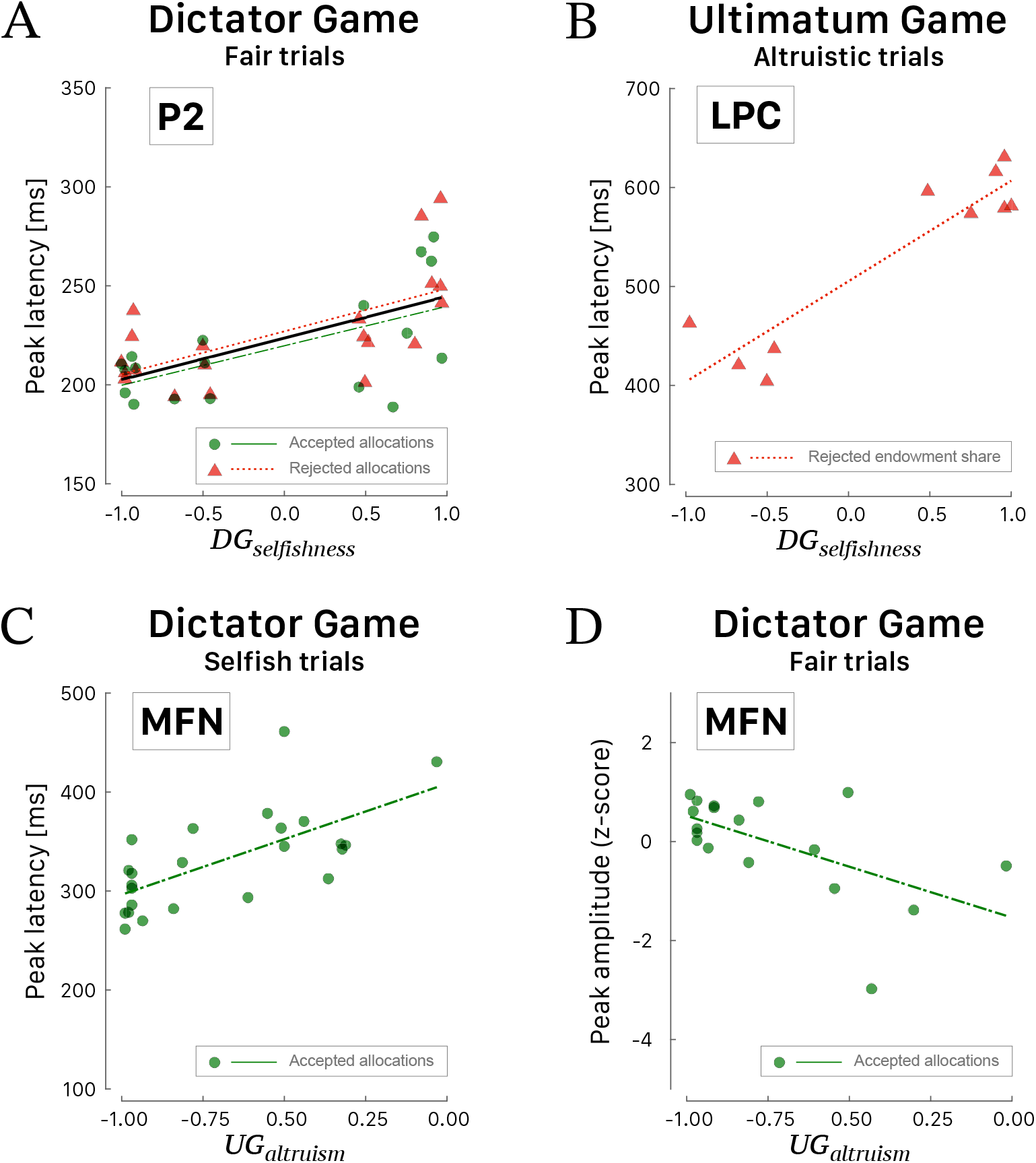
Scattergrams of significant correlations between ERP peaks and behavioral indices. **A.** Latency of P2 peak vs. *DG_selfishness_* during fair trials of DG. The black curve corresponds to the linear fit irrespective of the response choice. **B.** Latency of LPC peak *vs. DG_selfishness_* during altruistic trials of UG. **C.** Latency of MFN peak *vs*. U*G_altruism_* during selfish trials of DG. **D.** Amplitude of MFN peak vs. *UG_altruism_* during fair trials of DG. Green dots correspond to participants’ peak latencies when they agreed with the proposed allocation and red triangles when they refused the allocation. Green (long dashed) and red curves correspond to linear fitted regressions.

A positive correlation was also observed during altruistic trials of UG (Figure 8B) between the values of selfishness and the posterior-parietal LPC peak latency (*r* = 0.92, 95% CI [0.69, 0.98], *t*(8) = 6.657, *p* < .001) when the participants rejected the proposed payout (*F*(1, 8) = 44.31, *p* < .001, *R*^2^ = .847, 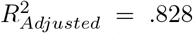). The only ERP wave that correlated well with altruism is the fronto-central MFN (Figure 8C,D) when the participants agreed with the proposed allocation. Interestingly, the smaller the altruism, the shorter the latency of MFN peak (*r* = 0.67, 95% CI [0.36, 0.84], *t*(22) = 4.186, *p* < .001) during selfish trials (*F*(1, 22) = 17.52, *p* < .001, *R*^2^ = .443, 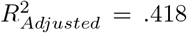) and the larger the altruism, the deeper the amplitude of MFN peak (*r* = −0.58, 95% CI [−0.82, −0.16], *t*(16) = −2.860, *p* < .001 during fair trials (*F*(1, 16) = 8.180, *p* = .011, *R*^2^ = .338, 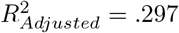).

## 4 Discussion

Human life is characterized by situations where the maximization of the individual payoff in face-to-face negotiations is challenged by the socio-cultural environment that contributed to shape moral phenomena such as altruism and fairness (Altman, 2005) and by the social distance to the other party, which affects the subjective perception of justice (Yu et al., 2015). In the context of an increasing number of contactless monetary transactions, our study aimed to test the hypothesis whether the characteristic patterns of brain activity—recorded by ERPs—were somatic markers associated with the behavioral profiles of individuals determined by a combination of Dictator and Ultimatum Game performances in the absence of effective human interactions between the parties. The analysis of the major ERP wave components showed that our observations were in agreement with several findings reported in the literature, which were mainly focused on only one of those classical neuroeconomic games (Nagel, 2001; De Martino et al., 2006; Fellner et al., 2009;

Güth, 2010; De Neys et al., 2011; Fiori et al., 2013). The identification of two homogeneous groups of participants, corresponding to selfish and fair participants, allowed to report more precisely the activity patterns associated with clearly modeled behaviors characterized by spiteful and costly punishments, which are respectively positively and negatively correlated with impulsive choices (Sanfey et al., 2003; Rilling and Sanfey, 2011; Rodrigues et al., 2018).

### 4.1 Behavioral results

Distinct groups of individuals were identified following a hierarchical cluster analysis based on indices associated with conceitedness/altruism and selfishness/fairness determined by performance during Ultimatum and Dictator Game. The definitions of conceitedness/altruism and selfishness/fairness are operational and we do not pretend that they necessarily overlap with semantic definitions that are culturally biased. The most representative and homogeneous groups were formed by fair participants (GrpF, individuals expressing a conceit behavior during UG and with a low level of selfishness during DG), by individuals characterized by less conceitedness and medium-high selfishness (GrpB), and selfish participants (GrpS, individuals with highest values of selfishness and lowest levels of altruism). The validity of the categorization of these participants’ groups is further confirmed by the observation that reaction times and, even more importantly, several characteristics of the ERP peaks —i.e., the somatic markers— were correlated with the behavioral indices. Hence, these indices are likely to reflect what they claim to reflect but they should not be extended beyond our experimental protocol without careful consideration.

In the UG, it is rationale to expect that the Proposer offers the smallest possible amount and the Responder accepts any offer. In this study, we could not fully evaluate a true altruistic attitude in the behavioral performance. All participants played the role of Responder and expressed conceitedness to some extent, because the most favorable payoffs for the Responder were equitable offers (i.e., with 40% or 50% of the share for the participant himself). The rejection of an unfair offer by the Responder in the UG is often explained by a bias towards profit maximization in association with positive social factors, such as common ethical principles and friendship, by negative factors, such as fear of the perceived consequences of having one’s offer rejected, and from the sense of guilt linked to worries about the opponent’s outcome (Carver and Miller, 2006; Marchetti et al., 2011; Gaertig et al., 2012). To test a Responder’s attitude from selfless/altruistic to truly greedy, even in the absence of a face-to-face interaction, UG’s task design would have to include endowment shares paying 60%–70% of the amount in favor of the Responder, as well as the most inequitable endowment shares (i.e., splits that provide 80%–90% of the amount to the Responder). We are aware of the limitations of the task design chosen in this study, which was driven by a compromise between the total duration of the experiment and the number of test repetitions that are necessary to get meaningful ERPs for both neuroeconomic games.

During DG, all participants played the role of of Allocator, who is the player who imposes an endowment share on the other party, and the behavioral performance distinguished two opposite groups, selfish (GrpS) and fair (GrpF) participants. However, during UG all participants played the role of Responder and both groups GrpS and GrpF tended to reject unequal offers (Table1). The rejection of the least favorable offers by the participants (i.e., payoffs of 10%-20% of the amount at stake) is considered to be the expression of a punishment towards a selfish Proposer (Forber and Smead, 2014; Ma and Hu, 2015) and was interpreted either as a spiteful punishment (Marlowe et al., 2011) or as costly —rather than altruistic— punishment (Brethel-Haurwitz et al., 2016; Yamagishi et al., 2017). The differences between the groups that during UG, when Responders have a passive position expecting to receive fair offers (Polezzi et al., 2008; Boksem and De Cremer, 2010), selfish participants (GrpS) were driven by spiteful punishment and fair participants (GrpF) by costly punishment. GrpA and GrpB participants also expressed a tendency towards selfishness, thus suggesting that spiteful behavior was predominant. This is not surprising because a single altruistic decision is likely to lack any reward for an individual, unless its repetition over time proves it is truly beneficial. The time frame of our experiment is very limited and the feature of learned behavior is in contrast to the spiteful punishment exercised by selfish participants who feel emotionally aroused when exposed to an unfair proposal even in a short lasting experimental context.

### 4.2 Socio-cultural dimension

Spiteful punishment is a kind of anti-social punishment applied in various ways that reflect the psychological and social dimensions of regional-cultural values (Chuah et al., 2007; Sylwester et al., 2013). It is recognized that the evolution of fairness is promoted by spitefulness and inhibited by altruism (Fehr and Gächter, 2002; Fehr and Rockenbach, 2004; Zhang and Fu, 2018). Behavioral differences have been observed in association with cross-country differences in socio-economic status and respect for authority (Inglehart, 2000; Oosterbeek et al., 2004; Chuah et al., 2007). High levels of cooperation and altruistic behavior can be observed in societies where people face economic disparities and poor social standing, and where they are exposed to punitive behavior if they violate certain norms and are treated as inappropriate (Fehr and Gächter, 2002; Nowak, 2006; Egas and Riedl, 2008).

In the present study, the socio-cultural dimension of our sample of participants is homogenous and made up of young Iranian males with an academic education. The choice to test participants without including both genders avoided the introduction of a gender effect discussed elsewhere in the literature (Eckel and Grossman, 2001; Servátka, 2009; García-Gallego et al., 2012; Li et al., 2018). Despite the evidence of altruistic tendencies in the Iranian people, such as willingness to pay for health services (Javan-Noughabi et al., 2017), organ donation (Abbasi et al., 2018), blood donation (Javadzadeh Shahshahani et al., 2006), moral sensitivity of nurses (Borhani et al., 2017; Amiri et al., 2020), the identification of the mechanisms associated with the perception of fairness has never been carefully investigated. Our results show that despite spitefulness appeared to prevail over fairness, we have observed a sizable sample of participants expressing costly/altruistic punishment and a tendency towards less conceit, somehow altruistic, behavior. It has also been established that in many experimental neuroeconomic contexts the game appears to reflect common interactional patterns of daily life, with greater behavioral variability between social groups than has been found among societies (Henrich et al., 2005), and the multiple play of UG does not affect rates of rejection (Oosterbeek et al., 2004). Therefore, we are confident that the participants included in this study did not introduce any strong behavioral bias towards an extreme behavioral strategy due to the homogeneous socio-demographic composition of our group of participants.

### 4.3 Fronto-central N1

In ERP studies, the latency and amplitude of a cortical wave component vary with the processing speed and the amount of cognitive resources required to reach the necessary degree of information processing for completion a certain mental task (Fabiani et al., 2000). The earliest ERP wave component that we have analyzed is the N1 negative component, much larger at fronto-central sites, in the 110–160 ms time window after trigger onset. It is important to distinguish the frontal N1 discussed here from the well known and described N1 component at occipital and temporal scalp sites associated with visual-spatial selective attention (Kutas et al., 1994; Mangun, 1995). The fronto-central N1 wave component is expected to be associated with a visual processing of a functional representation of the stimulus (Antal et al., 2000; Proverbio et al., 2007) and when participants expected a stimulus for decision making on whether to act or not to act (Di Russo et al., 2019). Such top-down prefrontal control of decision making (Fuster, 2015) suggests that N1 reflects a proactive cognitive control to access the procedural knowledge necessary to enable an active motor response (Petit et al., 2006; Proverbio et al., 2007; Di Russo et al., 2019).

We observed that fronto-central N1 did not vary much between the games, but it was strongly associated with the group of participants. The greater the selfishness, the shorter the latency of N1 and in the group of selfish participants this component peaked about 10 ms earlier than fair participants and with smaller amplitude. In our tasks, the visual cues carry a cognitive value associated with the costs and benefits determined by the behavioral/personality traits of each individual. Hence, we can interpret the N1 difference between fair and selfish participants as the first evidence in ERPs between participants who have otherwise expressed costly and spiteful forms of punishment. Longer N1 latencies and larger amplitudes in fair participants are compatible with a top-down control of a rather rich and less stereotyped set of initial activity patterns. Our results might also be interpreted in agreement with an anterior N1 enhancement assumed to reflect the cost of a modality change and readjustment of attentional weight-setting in order to optimize target detection (Töllner et al., 2009; Hsieh and Wu, 2011).

### 4.4 Fronto-central P2

The P2 wave peaked in the range of 180 to 260 ms and was most salient in the fronto-central area, where a top-down process using previous experience and expectations is indicative of stimulus evaluation rather than response production (Potts, 2004; Rauss and Pourtois, 2013; Fuster, 2015). Increased fronto-central P2 is expected in response to rewarding contextual outcomes, in participants who feel gratification following the amount of cognitive resources activated by a reward signal in the brain, which may include satisfaction when observing the opponent’s misfortune (Falco et al., 2019; Weiss et al., 2020). In previous decision-making studies, it was also suggested that P2 is modulated by the degree of predictability of the outcomes (Polezzi et al., 2008) and by the social context (Hu et al., 2014; Liu and Huang, 2015). Our results are in agreement with the rationale that refusal of fair payoffs is a rewarding choice for selfish participants. and refusal of selfish payoffs is a rewarding choice for fair participants. These findings support the hypothesis that in DG a selfish player required more cognitive resources when he refused to allocate a fair share and a fair player required more cognitive resources when he agreed to assign a selfish profit. In UG, the analysis of P2 wave suggested that selfish participants tend to maximize their gain with greater and longer P2 even after acceptance of less favorable payoffs. Moreover, we found evidence that accepting less favorable payoffs was a rewarding choice for the group of more altruistic participants (or less conceit, GrpB).

### 4.5 Medial Frontal Negativity

The MFN is a wave component that has been observed in association with some assessment of fairness (Botvinick et al., 1999; Gehring and Willoughby, 2002; Alexopoulos et al., 2012; Li et al., 2020), social status and altruistic bias (Van der Veen and Sahibdin, 2011; Sun et al., 2015; Cui et al., 2019; Mayer et al., 2019), but only one study has reported quantitative ERP results in Allocators during DG (Li et al., 2020). However, these studies did not clearly distinguish between behavioral groups. In our study, we observed that in both games the negativity of MFN tended to be greater when the latency was shorter. In general, the greater the selfishness, the deeper the negativity of MFN. Being associated with disadvantageous inequitable offers and with altruistic tendency, MFN is expected to be larger in altruistic participants. This was indeed our observation for GrpB participants playing the Dictator Game, who expressed less conceit and more altruistic behavior. This observation was further confirmed by the correlation of MFN amplitude and the index *UG_altruism_* in the fair trials of DG when the participants playing the role of Allocator accepted the proposed amount.

In fair participants, the latency of MFN was shorter during DG after accepting to allocate a fair amount to the other player than after rejection of a selfish allocation, and during UG the latency of MFN was longer after acceptance of most advantageous over acceptance of least favorable splits of the endowment share. In selfish participants, spiteful punishment was revealed by shorter latency and reduced MFN negativity after rejection of the least favorable payoffs over acceptance of most advantageous offers. These results support the hypothesis that in UG the MFN amplitude of Responders is likely to be associated with an emotional reaction to the expectation of fairness by the Proposers. Thus, the perception of the offers by the Responders —all participants of this study played this role— depended on the behavioral group, and not on the absolute value of the endowment share (Van der Veen and Sahibdin, 2011; Mayer et al., 2019; Spapé et al., 2019; Li et al., 2020). In general, our results may be interpreted in accordance with the hypothesis of a short latency of MFN as a sign of an emotional response, which required little cognitive evaluation and expressed reckless mental processing (Brookhuis et al., 1981; Kok, 1997; Codispoti et al., 2007; Fukushima and Hiraki, 2009; Boksem et al., 2011). This is also in agreement with the presumed generation of MFN in the anterior cingular cortex in association with emotional regulation (Etkin et al., 2011; Stevens et al., 2011; Kanske and Kotz, 2011; García-Cabezas and Barbas, 2017).

### 4.6 Posterior-parietal late positive component

Evidence reported in the literature indicates that LPC-associated activity is likely to be generated by a widespread network of cerebral sources including the frontal lobe, the cingulate cortex and the parietal regions involved in attention control (Sabatinelli et al., 2007; McDonald and Green, 2008; Scharmüller et al., 2011; Sun et al., 2017). LPC is closely related to relevant motivational stimuli, to the subsequent sustained allocation of attention (Ito et al., 1998; Wu et al., 2011b; Bamford et al., 2015), and to the saliency of an outcome also in social comparisons (Sun et al., 2020; Zhang et al., 2021; Sharif et al., 2021). It could be hypothesized that a large LPC wave should characterize selfish Allocators playing the Dictator Game. In both games, we have indeed observed greater LPC (and at shorter latency in UG) in selfish over fair participants. Selfish and fair participants were characterized by quiet opposite patterns of activity recorded at the posterior-parietal area. Recent evidence provided by the analysis of a modified Dictator Game exploring the role of players’ social status (Cui et al., 2019) revealed that LPC amplitude was more sensitive to the interaction of the recipient’s status and fairness— i.e., with a tendency of greater amplitude triggered by unfair endowment shares for a high-status Recipient and greater amplitude triggered by fair splits for a medium-status Recipient. It is interesting to note that selfish participants were also characterized during UG by a sustained late positive wave at the unusual fronto-central sites. This observation suggests that in these participants a potential source probably not located at the cortical level, but deeper in the brain, might contribute to a late positive wave visible at fronto-central as well as at posterior-parietal areas.

It is rationale that altruistic players expect equitable shares of the endowment during the Ultimatum Game. Being associated with motivation and subsequent allocation of attention, the amplitude of posterior-parietal LPC is expected to be grater in altruistic Responders playing the Ultimatum Game. In agreement with this hypothesis, we observed greater amplitude of LPC in GrpB participants when they accepted equitable over inequitable splits of the sum at stake. Moreover, during DG the latency of LPC was shorter in GrpB Allocators when they refused more advantageous allocations over refusal of fair allocations. Overall, the differences between selfish and fair participants support the hypothesis that LPC at posterior-parietal sites reflects a processing stage that evaluates norms of fairness (Wu et al., 2011b) in association with saliency interacting with context (Minnix et al., 2013; Stolz et al., 2019; Sun et al., 2020; Bauer et al., 2021).

### 4.7 Limitations

A critical limitation of our study is common to most neuroeconomic studies, that is whether studies based on experimental protocols in laboratory settings can really give insights in human brain activity outside real-world situations (Clithero et al., 2008; Huettel and Kranton, 2012; Konovalov and Krajbich, 2019). There is no doubt that interaction between real people plays a key role in business relationships, which are based on a much more complex set of contextual signals with an individual realm of emotions and feelings that help shape individual personalities. In our protocol, the participants were informed that they were playing against a human party located elsewhere, without specifying whether the other party was the experimenter or other participants. This is a workaround, but it is important that the participants were not simply playing for nothing because they ultimately received a real amount of money that depended on the performance accumulated at the games. Future studies should also require the participants to fill questionnaires defining personality traits (e.g., HEXACO; Ashton et al., 2014) but a validated Persian version of this inventory scale (Basharpoor et al., 2019) was not yet avaibale at the time our study was run and completed.

The semantic definition of benefit in the economic context set by neuroeconomic games refers to a rational choice aimed at achieving the optimal maximization of gain (Bernheim, 2009). The definitions of altruism, selfishness and fairness are very subjective, but we have defined the sets of conceit/altruistic and selfish/fair trials within a precise operational context defined by our experimental protocol, based on the same participant playing both games. The observation of irrational choices, i.e. low levels of altruism and opposing levels of selfishness (GrpF and GrpS), indicate that one’s own assessment of the coherence of actions can lead to beneficial outcomes that are biased compared to a rational evaluation of monetary payment. To this respect, another limitation of the current study is the lack of neurophenomenological interview (Bockelman et al., 2013; Høffding et al., 2021; Klinke and Fernandez, 2022) in order to understand the subject’s own perception of his/her spiteful or costly punishment.

Last but not least, we recognize methodological limitations associated with the sample size when we consider that the number of trial repetitions needed for averaging ERPs was limited by the duration of one session and the unbalanced number per set determined by individual strategies that could not be foreseen before the experiment. We are aware that peak latencies and amplitudes reported in our study were sensitive to high-frequency noise. The signal-to-noise level is low in our study due to averaging over a small amount of trials and this brings some methodological considerations. The limits of the time windows of an ERP wave are extremely difficult to set firmly for all participants because of the individual differences in the adopted strategy and may vary between participants across the various sets of trials (selfish trials: accepted vs. rejected, fair trials: accepted/rejected). These time limits could be considered only after a careful post-hoc analysis, which is contradictory with the measurement of the mean amplitude (instead of peak amplitude) and fractional area latency (instead of peak latency) that are usually considered more meaningful values in itself (Fabiani et al., 2007; Woodman, 2010; Luck, 2014). Our aim was not the establishment of standardized ERP values for the experimental protocols of neuroeconomic games, but to determine if somatic markers could be reliably observed in participants who clearly adopted different strategies. In the absence of a large number of trials at hand to overcome the difficulties in measuring peak values with a jackknife-based approach (Kiesel et al., 2008), we have adopted the method of repeated measurements taken by several observers. In future studies, we believe that the newly presented standardized measurement error for amplitude and latency values carries the potential for greater replicability of ERP studies in neuroeconomic experimental protocols, making it possible to identify participants with low-quality data and noisy channels (Luck et al., 2021).

### 4.8 Conclusion

This study has presented a novel combination of Dictator and Ultimatum Game with the aim of investigating whether ERP markers support the hypothesis that spiteful and costly punishments can be differentiated. We have presented evidence that different groups of participants can be identified on the basis of their performance at the combination of DG and UG. Two groups of participants, both with little tendency towards altruism, were characterized by a selfish or fair attitude during Dictator Game. An early ERP negative component (N1), peaking in the interval of 110 to 160 ms at fronto-central sites, distinguished fair participants from selfish ones. Analysis of middle frontal negativity (MFN) suggested that selfish participants were likely to exhibit spiteful punishment positively related to impulsive choice. The differences in late positive component (LPC) observed between the groups could be interpreted as that costly punishment behavior, enabled by fair participants, must be stable to make sense and requires less attention than selfish participants, whose cognitive-behavioral contradiction requires more attention and activation of a larger brain network to complete the information processing necessary to make a decision. In conclusion, this study brings an original contribution and new evidence to the existence of different circuits activated by the evaluation of fair and unfair proposals in participants characterized by different expressions of perception of fairness.

## Author Contributions

Conceptualization, AMM, HP, MAM, RK; formal analysis, HP, MAM, AL; funding acquisition, HP, AL; clinical investigation, MAM; methodology, AMM, HP, AL; project administration, MAM, HP, AL; supervision, HP, AL, AEPV; validation, HP, AL, AEPV, AL; visualization, AEPV, AL; writing original draft, AMM, AEPV, AL; writing-review and editing, AEPV, AL.

## Acknowledgments

The authors extend their gratitude and acknowledgements to all study participants and study team members for their time and energy spent on this project.

